# Hypomethylating agents induce epigenetic and transcriptional heterogeneity with implications for AML cell self-renewal

**DOI:** 10.1101/2024.01.30.577864

**Authors:** Danielle R Bond, Sean M Burnard, Kumar Uddipto, Kooper V Hunt, Brooke M Harvey, Luiza Steffens Reinhardt, Charley Lawlor-O’Neill, Ellise A Roper, Sam Humphries, Heather C. Murray, Abdul Mannan, Matthew D Dun, Charles E de Bock, Nikola A Bowden, Anoop K Enjeti, Nicole M Verrills, Carlos Riveros, Kim-Anh Lê Cao, Heather J Lee

## Abstract

DNA hypomethylating agents (HMAs) are used to treat acute myeloid leukemia (AML) and myelodysplasia patients who are unsuitable for intensive chemotherapy. However, low response rates and therapy-resistant relapse remain significant challenges. To improve outcomes, we must understand how AML cells survive HMA treatment and continue to proliferate following therapy. We combine single-cell multiomics with parallel colony-forming assays to link HMA-induced heterogeneity with functional consequences in AML cellss. Azacytidine (AZA) and decitabine (DAC) induced global epigenetic heterogeneity, associated with upregulation of inflammatory responses and cell death pathways in a subset of hypomethylated cells. Some cells maintained high DNA methylation levels during treatment, and these methylation-retaining cells had increased self-renewal capacity following DAC treatment in two FLT3-ITD AML cell lines. Transcriptional profiling of colonies formed after HMA treatment revealed many genes with altered expression in both methylation-retaining and hypomethylated cells, with increased expression of cholesterol-related genes observed in all cell lines. Inhibition of the cholesterol biosynthesis pathway by rosuvastatin enhanced HMA effects on colony formation in vitro and extended survival in two in vivo models of AML. Our study demonstrates that HMA-induced epigenetic heterogeneity has implications for AML cell growth and identifies statins as a candidate co-treatment strategy to improve HMA efficacy.

## Introduction

Hypomethylating agents (HMAs), such as azacytidine (AZA, 5-azacytidine) and decitabine (DAC, 2’-deoxy-5-azacytidine), are used to treat patients with acute myeloid leukemia (AML) and myelodysplastic syndrome (MDS) (1). AZA and DAC are cytidine analogs, which are incorporated into DNA during replication (1, 2). This leads to the degradation of DNA methyltransferase (DNMT) enzymes (3) and loss of DNA methylation in subsequent cell divisions (2, 4). HMA mechanisms of action have been difficult to precisely define because genome-wide loss of methylation is associated with pleiotropic transcriptional changes (5). HMA-induced promoter hypomethylation permits the re-expression of tumor suppressor genes (6), re-activation of DNA repair pathways (7) and upregulation of differentiation markers in AML cells (8). HMA treatment can also increase the expression of cancer testis antigens (9) and enhance antigen presentation in cancer cells (10). Furthermore, genome-wide de-repression of transposable elements (TEs) has been shown to trigger a viral mimicry response, in which dsRNA stimulates interferon signaling and apoptosis (11–13). Unlike DAC, AZA can also be incorporated into RNA, which influences transcript stability and translation (14, 15).

While single-agent HMA treatment extends survival in many patients (16–18), only 20% show a complete response (CR) to therapy (19). Responses are also short-lived (e.g., 8-15 months (20)), with most patients developing HMA-resistant relapse. To address these limitations, clinical trials have tested co-treatment strategies, with some delivering improved outcomes (5). For example, the pro-apoptotic therapy venetoclax has increased response rates in elderly AML patients undergoing HMA treatment (21). However, relapse remains a significant problem, with the median duration of response being 11-18 months for patients treated with both venetoclax and HMA therapy (21, 22). To improve the long-term benefits of HMA therapy, it is essential to understand how AML cells survive treatment and proliferate to cause relapse.

In this study, we characterize HMA-induced molecular heterogeneity with implications for the self-renewal capacity of AML cells. We perform single-cell multiomic analysis to reveal global HMA-induced DNA methylation heterogeneity and methylation-retaining cells that appear to evade treatment. In parallel colony-forming assays, we show that methylation-retaining cells have a growth and self-renewal advantage following treatment with DAC, but not AZA, in two FLT3-ITD cell lines.

We also identify the upregulation of cholesterol-related genes in colonies formed after HMA treatment, and identify statins as a candidate treatment strategy to enhance the long-term efficacy of HMAs.

## Results

### DNA methylation heterogeneity induced by HMA treatment

We performed single-cell analysis of DNA methylation in AML cell lines (HL-60, MOLM-13, and MV-4-11; Supplementary Table 1) treated with low doses of decitabine (DAC) or azacytidine (AZA) to induce maximal demethylation with minimal effects on cell growth and viability (Supplementary Fig. S1). After three days of HMA treatment, a striking level of DNA methylation heterogeneity was observed (Fig. 1A; Supplementary Table 2). While untreated cells showed homogeneously high levels of DNA methylation (e.g., HL-60: 65-73%), the extent of hypomethylation varied substantially among cells treated with DAC (e.g., HL-60: 17-69%) or AZA (e.g., HL-60: 20-69%). Interestingly, a small proportion (1-5%) of methylation-retaining cells (Fig. 1A, dashed boxes) displayed no evidence of HMA-induced hypomethylation. The extent of HMA-induced hypomethylation was related to cell division rate, as indicated by positive correlations between DNA methylation and CellTrace fluorescence (Fig. 1B; Supplementary Fig. S2). This is consistent with HMA incorporation during replication (2) and confirms that slowly dividing cells can avoid the effects of HMA treatment.

**Figure 1.**
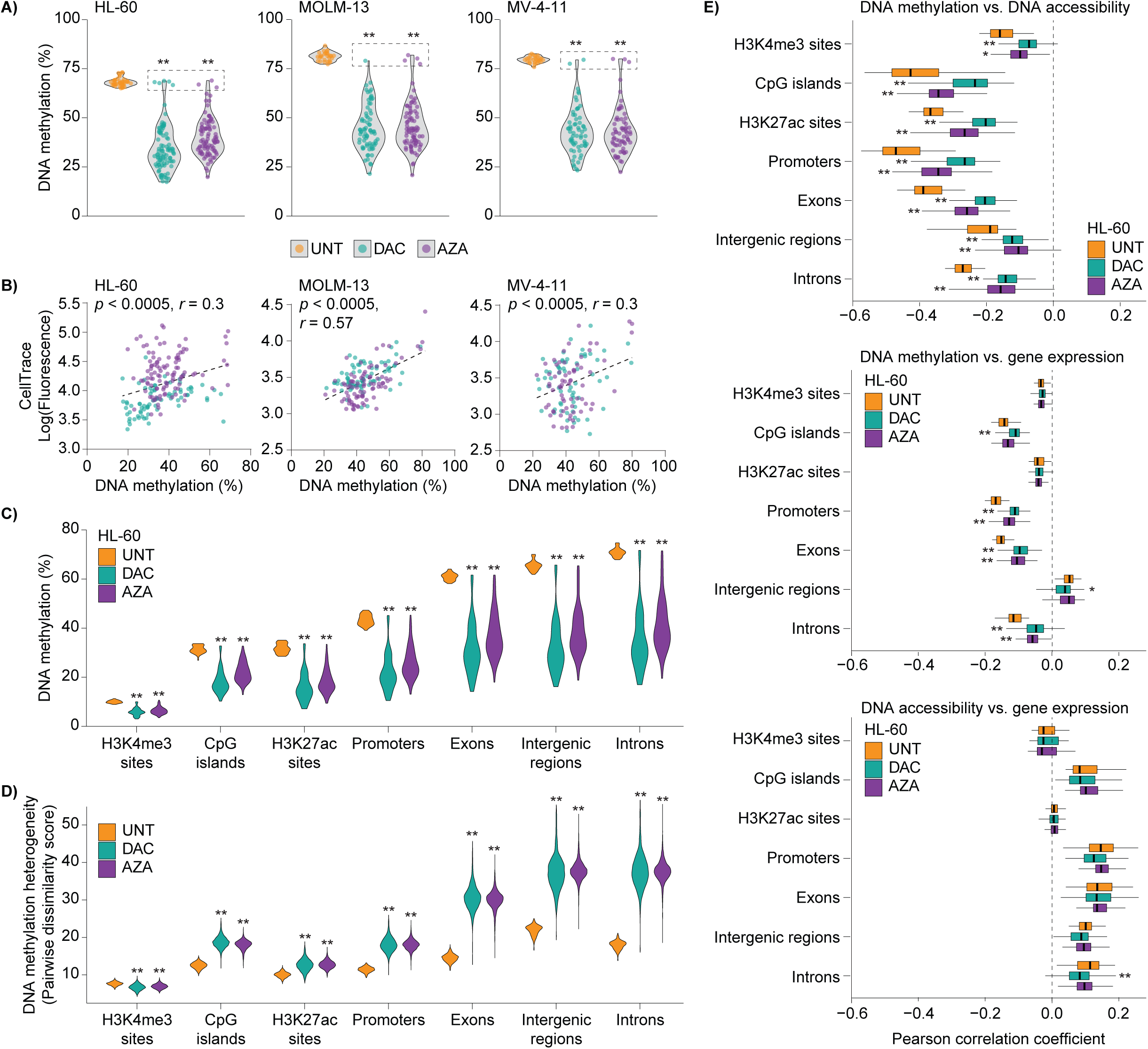
HMA treatment induces DNA methylation heterogeneity in AML cells. HL-60, MOLM-13 and MV-4-11 cells were labelled with CellTrace and treated with decitabine (DAC; 100nM) or azacytidine (AZA; HL-60: 2000nM, MOLM-13 and MV-4-11: 500nM) every 24h for 72h. Single cells collected by indexed FACS were subjected to scNMT-seq (HL-60) or scTEM-seq (MOLM-13, MV-4-11). A) Violin plots of DNA methylation levels in single HL-60 (left), MOLM-13 (middle) and MV-4-11 (right) cells. Superimposed points show single-cell values from untreated (UNT, orange), DAC (cyan) and AZA (purple) groups. Dashed boxes surround DAC and AZA cells with methylation levels within the range of UNT samples. Data are shown for 185-222 cells collected from 2-3 replicate experiments in each cell line (UNT, n = 27-38; DAC n = 63-93; AZA n = 68-91). B) Scatter plots comparing CellTrace fluorescence and DNA methylation in single cells, with linear regressions, F-test p-values, and Pearson correlation coefficients (r). C) Violin plots of DNA methylation levels in different genomic contexts from HL-60 scNMT-seq data. D) Violin plots of DNA methylation heterogeneity, as determined by pairwise dissimilarity analysis, within different genomic contexts from HL-60 scNMT-seq data. E) Box and whisker plots of Pearson correlation coefficients computed within single cells from HL-60 scNMT-seq data. DNA methylation and DNA accessibility were considered in different genomic contexts, and individual loci were matched based on genomic co-ordinates. Correlations were performed between DNA methylation and DNA accessibility (top), DNA methylation and gene expression (middle), and DNA accessibility and gene expression (bottom). Boxes depict the interquartile range (IQR) with median. Whiskers extend to the highest and lowest data points within 1.5 x IQR of the first and third quartile. Outlying data points are not shown. Statistical analysis was performed using ordinary one-way ANOVA with Dunnett’s (A) or Šídák’s (C-E) multiple comparisons test: * p < 0.05, ** p < 0.0005 vs. UNT.

Genomic contexts with high levels of DNA methylation in untreated HL-60 cells (such as exons, introns, and intergenic regions) showed the most pronounced reduction following HMA treatment (Fig. 1C), with this loss of methylation accompanied by increased cell-to-cell heterogeneity (Fig. 1D). In contrast, active promoters marked by H3K4me3 were the only genomic features to show a significantly decreased DNA methylation heterogeneity after HMA treatment, likely due to their low levels of DNA methylation in untreated cells.

To explore links between DNA methylation heterogeneity and other layers of genetic regulation, we used multi-omic data collected from HL-60 cells. The scNMT-seq method profiles DNA methylation, DNA accessibility, and gene expression in parallel (23), allowing these molecular modalities to be correlated with each other across the genome of individual cells (Fig. 1E). Untreated cells showed the expected trends, with DNA methylation being negatively correlated with accessibility and gene expression, whereas DNA accessibility and transcription were positively correlated. In HMA-treated cells, the relationship between DNA methylation and accessibility was significantly weakened in all genomic contexts. HMA treatment also significantly weakened the correlation between DNA methylation (in promoters, exons, and introns) and expression of associated genes. In contrast, HMA treatment had a minimal effect on the association between DNA accessibility and gene expression. These observations imply that loss of DNA methylation is not always accompanied by increased accessibility and transcription.

### Transcriptional programs linked to HMA-induced epigenetic heterogeneity

To clarify how HMA-induced epigenetic heterogeneity influences transcriptional responses, we performed a multivariate analysis to integrate the three molecular layers: DNA methylation, DNA accessibility, and transcription. We excluded untreated cells to focus on variability among HMA-treated cells and applied an unsupervised sparse Partial Least Squares (sPLS) method (27). This approach was used to identify variably expressed transcripts that are highly correlated to changes in DNA methylation and accessibility in regulatory regions (CpG islands, promoters, H3K4me3 sites, and H3K27ac sites) and 3kb genomic windows.

The 200 transcript features selected by sPLS were further examined in both treated and untreated cells, revealing 4 groups of cells with distinct transcriptional profiles (Fig. 2A, B). A significant and negative correlation (r = −0.37, p < 1 x 10^-7^) between TE LINE:L2a expression and component 1 was observed (Fig. 2C; Supplementary Fig. S3A), consistent with activation of viral mimicry in cell group 3. However, there was no significant relationship between global DNA methylation and component 1, and cell group 3 had similar methylation levels to other HMA-treated cells (Fig. 2D; Supplementary Fig. S3D). This indicates that global methylation levels are insufficient to explain the transcriptional responses of HMA-treated AML cells.

**Figure 2.**
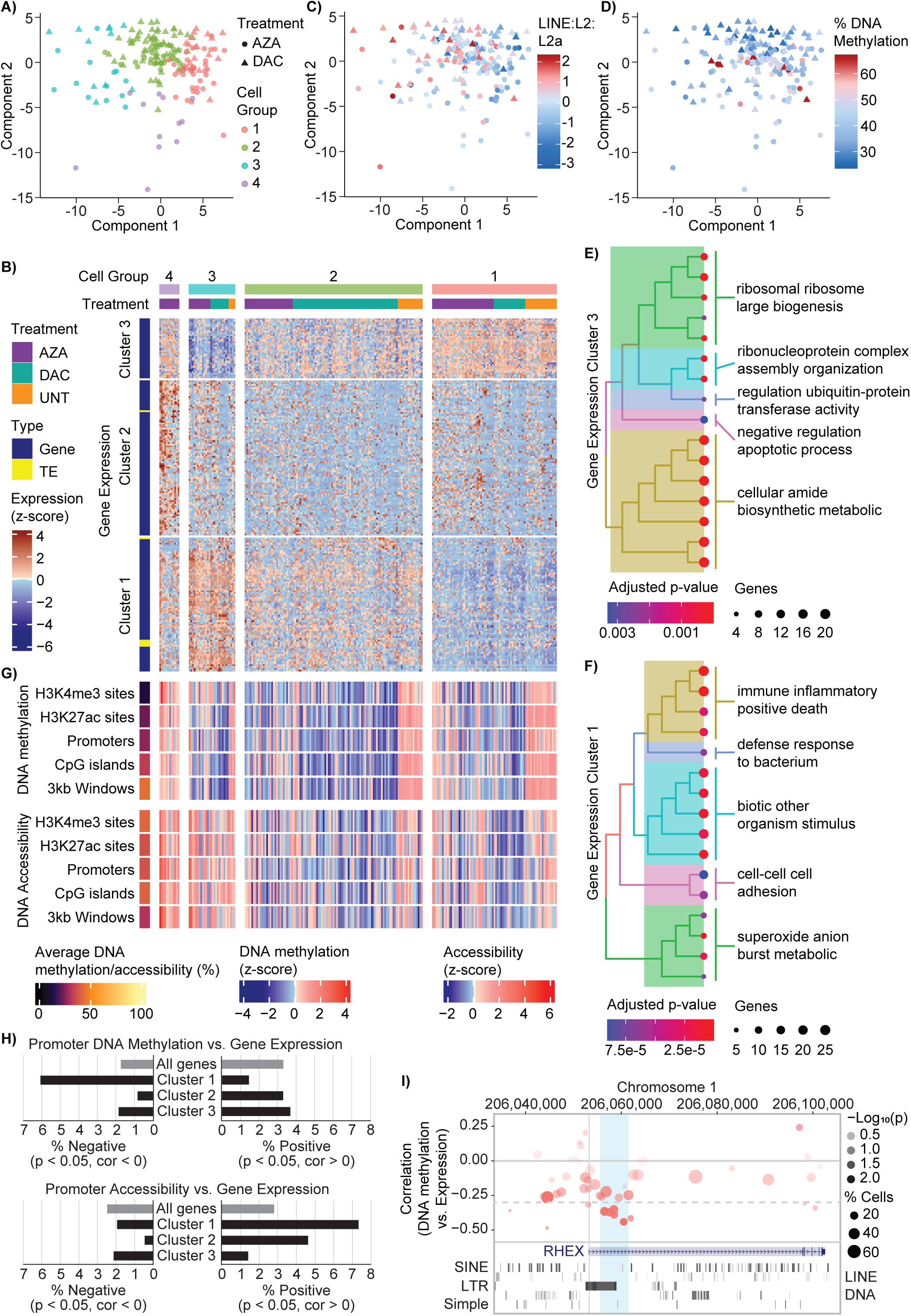
HMA-induced epigenetic heterogeneity influences transcriptional programs in AML cells. HMA treated (AZA and DAC only) HL-60 scNMT-seq data underwent multivariate feature selection by sparse least squares (sPLS) using an unsupervised model. A) sPLS projection of cells based on transcript features coloured by cell group 1-4 (from B). B) Heatmap of transcript features selected by sPLS displaying all samples (treated and untreated) as columns, are split by k-means clustering and grouped by treatment. Individual gene and TE expression levels (rows) are z-score normalised and split by k-means clustering with internal hierarchical clustering. C-D) sPLS projections of cells based on transcript features are coloured by C) LINE:L2:L2a expression, and D) global methylation level. E-F) Tree plots of the gene ontology (GO) over-representation analysis (ORA) for gene expression clusters 3 and 1. G) Heatmaps summarising the DNA methylation and DNA accessibility features selected by sPLS. Samples (columns) are ordered according to the heatmap in B. The average methylation or accessibility of all sPLS selected features across all samples (treated and untreated) is displayed on the left for each genomic context. The two heatmaps show z-score normalised averages of DNA methylation and accessibility for all sPLS selected features in each genomic context. H) Pearson correlations were computed between gene expression and DNA methylation (top) or accessibility (bottom) of associated promoters. Bar graphs show the percentage of correlations (p < 0.05) with negative (left) and positive (right) coefficients for all genes and filtered by gene expression cluster (identified in D). I) Correlations between RHEX expression and DNA methylation of adjacent loci (top) are displayed together with annotated short and long interspersed nuclear elements (SINE, LINE), long terminal repeat (LTR), DNA, and Simple repeat sequences (bottom). The blue shading highlights a promoter-proximal region of intron 1 that included 6 regions with negative correlations (cor < −0.3, p < 0.05) between DNA methylation and RHEX expression.

Gene ontology (GO) over-representation analysis (ORA) was performed for each of the three clusters of transcript features (Fig. 2B; Supplementary Table 3). Cell group 3 had low expression of genes in cluster 3, which were related to translation and inhibition of cell death (Fig. 2E; Supplementary Table 4). Cell group 3 also had high expression of genes in cluster 1, which were enriched in terms related to immune inflammatory response and positive regulation of cell death (Fig. 2F; Supplementary Table 4). This transcriptional profile is consistent with the expected effects of HMA treatment (1), and 29 of the 78 genes in cluster 1 were significantly upregulated by DAC and/or AZA in matched bulk RNA sequencing (RNA-seq) data (Supplementary Table 3). In contrast, cell group 1 displayed an inverted gene expression pattern and included many untreated cells (observed:expected ratio = 1.48). This suggests that cell group 1 did not activate transcriptional pathways commonly associated with HMA treatment, despite low methylation levels in most cells (Fig. 2A, D). The expression profile of cell group 2 was intermediate between groups 1 and 3, whereas cell group 4 showed distinctive upregulation of genes expressed in cluster 2. No ontology terms from cluster 2 retained significance after multiple testing correction.

The sPLS model also selected epigenetic features that were correlated to the 200 transcript features, without considering the direction of correlation nor the proximity of features (Tables S5, S6). For cell group 3, the selected DNA methylation features have low methylation in many cells, whereas the DNA accessibility features tend to have high accessibility (Fig. 2B, G). This was in contrast to other cell groups, suggesting that the transcriptional response to HMA treatment is associated with reductions in DNA methylation and gains in accessibility. Consistently, transcript features from expression cluster 1 had predominantly positive correlations with accessibility features and many negative correlations with methylation features, especially in CpG islands and 3kb genomic windows (Supplementary Fig. S4).

To test whether epigenetic alterations in cis-regulatory elements could influence transcriptional responses to HMA treatment, we next correlated gene expression with DNA methylation and accessibility in nearby genomic loci. Genes from expression cluster 1 showed a significant shift toward negative correlations with promoter methylation (p = 2.15×10^-11^, χ^2^ test) and positive correlations with promoter accessibility (p = 2.26×10^-8^, χ^2^ test), suggesting that these genes are particularly sensitive to loss of DNA methylation in cis (Fig. 2H, I; Supplementary Table 7). An interesting example is RHEX (regulator of hemoglobinization and erythroid cell expansion), which is highly expressed in AML (32). Negative correlations between RHEX expression and DNA methylation were concentrated in a promoter-proximal region of intron 1, which contains several conserved long terminal repeat (LTR) TEs (Fig. 2I).

In summary, our single-cell multiomic analysis identified transcriptional changes linked to patterns of epigenetic heterogeneity in HMA-treated AML cells. Importantly, activation of genes involved in inflammatory response and cell death pathways (expression cluster 1) was observed in only a subset of hypomethylated cells (cell group 3) (Fig. 2A, C, D).

### Effects of HMA-induced heterogeneity on the self-renewal capacity of AML cells

To determine the consequences of epigenetic and transcriptional heterogeneity on AML cell self-renewal, colony-forming assays were performed. DAC and AZA significantly decreased colony counts (Fig. 3A), as previously reported (33). DAC treatment also significantly increased the size of colonies in the two FLT3-ITD cell lines, MOLM-13 and MV-4-11 (Supplementary Table 1, Fig. S5A). To characterize the molecular profiles of colonies formed after HMA treatment, we performed single-colony analysis of DNA methylation (Fig. 3B-C; Supplementary Table 8). Following DAC treatment (Fig. 3B, solid fill) many colonies (34-65%) had DNA methylation levels comparable to untreated colonies (≥75%). This was in stark contrast to the low percentage of methylation-retaining cells (1-5%) observed in single-cell data after 72h of DAC treatment (Fig. 1A, dashed box; Fig. 3B, dashed line). Surprisingly, this shift toward higher DNA methylation levels was far less pronounced in colonies established following AZA treatment (Fig. 3C).

**Figure 3.**
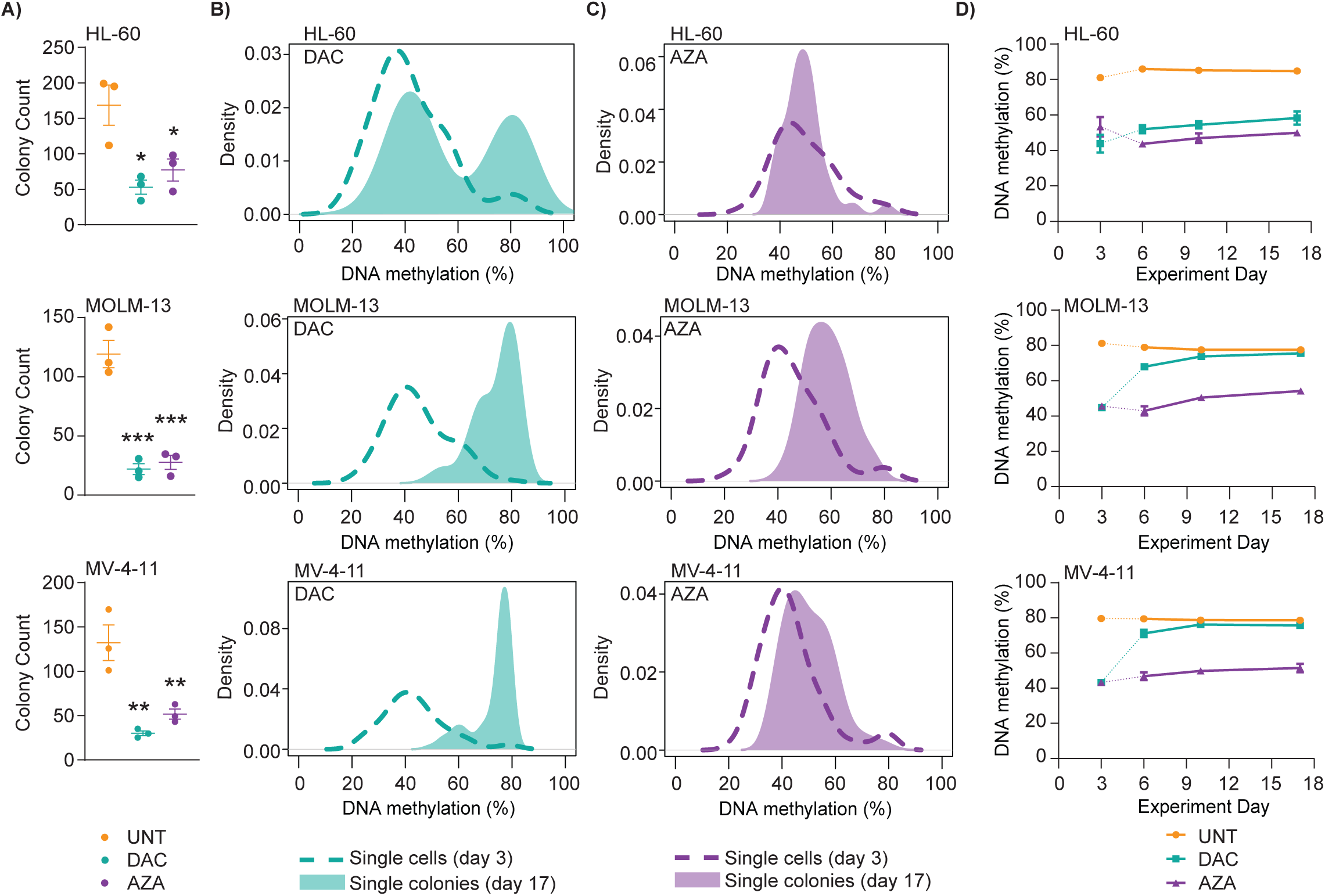
Highly methylated AML cells display a growth and survival advantage following treatment with DAC, but not AZA. A) HL-60, MOLM-13, and MV-4-11 colony counts following treatment with DAC (cyan) or AZA (purple) vs. untreated cells (UNT, orange). B-C) Density plots show the average DNA methylation levels for single cells collected on experiment day 3 (dashed line) and individual colonies collected on experiment day 17 (solid fill) following treatment with DAC or AZA. HL-60 scNMT-seq data were filtered for cytosines within SINE Alu sites for direct comparison to scTEM-seq data from colonies. Data are shown for 288 colonies collected from triplicate experiments in each cell line (n = 96 per treatment). D) Time-course experiment showing changes in average DNA methylation of cells collected at different time points throughout the colony-forming assay (experiment days 6, 10 and 17). Values for experiment day 3 were obtained from single-cell data (Fig. 1A). Data in A and D are expressed as mean +/-standard error of the mean (SEM). Statistical analysis of colony counts (A) was performed using ordinary one-way ANOVA with Dunnett’s multiple comparisons test with a p < 0.05 cut-off for significance (p < 0.03*, p < 0.006**, p < 0.0002***).

To assess the recovery of DNA methylation following HMA treatment, time-course analysis was performed during colony formation (Fig. 3D). DNA methylation levels in HMA-treated HL-60 colonies were low throughout the time course (Fig. 3D, top), mirroring an analysis performed in suspension culture (Supplementary Fig. S1E). Similar results were obtained for MOLM-13 and MV-4-11 colonies derived after AZA treatment (Fig. 3D, center and bottom). In contrast, high levels of DNA methylation were observed in the early stages of colony formation (experiment day 6) following DAC treatment of MOLM-13 and MV-4-11 cells (Fig. 3D, center and bottom). This suggests that the shift toward high DNA methylation observed in these colonies (Fig. 3B) was not due to gradual recovery of methylation. Rather, our data are consistent with the selection of highly methylated cells in colony-forming assays performed after DAC treatment. We deduce that methylation-retaining cells can have increased self-renewal and proliferative capacity relative to hypomethylated cells after HMA treatment, but this may depend on the mutational profile of the AML cells and the HMA used.

### Transcriptional changes in hypomethylated and methylation-retaining colonies

Next, we generated matched transcriptomes from the same set of colony samples (Supplementary Table 8). Single-colony RNA-seq and principal component analysis (PCA) showed that DAC and AZA samples were generally distinct from untreated samples, regardless of their global DNA methylation levels, in all cell lines (Fig. 4A). This implies that HMA exposure has substantial effects on the transcriptome, even in highly methylated cells.

**Figure 4.**
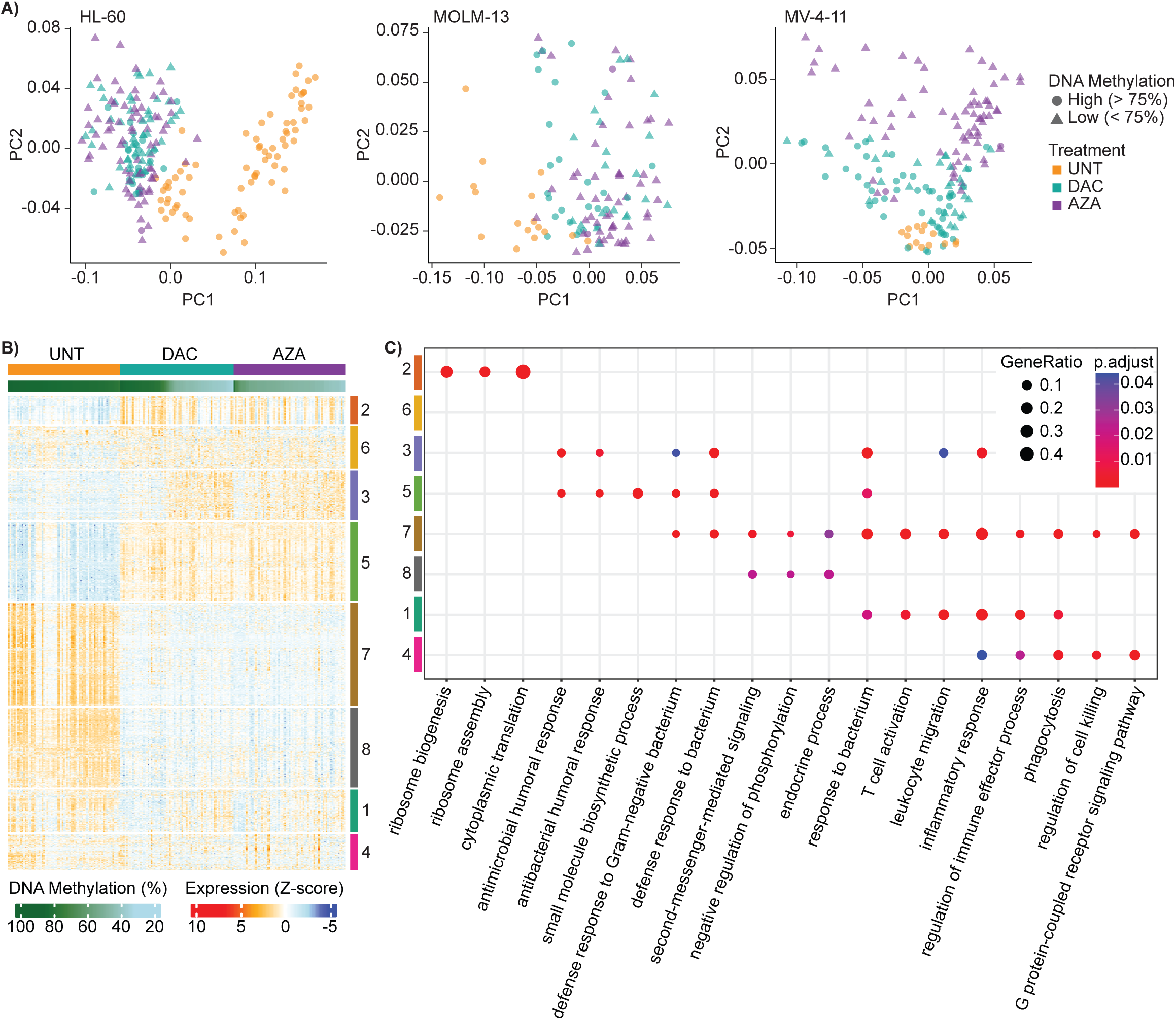
HMA treatment has sustained transcriptional effects in both methylation-retaining and hypomethylated cells. A) Principal Component Analysis (PCA) plots of single-colony RNA-seq data from AML cell lines, highlighting treatment groups (UNT = orange; DAC = cyan; AZA = purple) and matched mean DNA methylation levels (circle: high > 75%; triangle: low < 75%). Data shown for 119 - 220 colonies collected from 3 replicate experiments in each cell line (UNT, n = 14-73; DAC, n = 46-78; AZA, n = 56-73). B) Heatmap of the top 2000 highly variable genes from colony RNA-seq data. Samples are ordered by decreasing global methylation levels (green gradient) within each treatment group. Rows are grouped by K-means clusters based on gene expression, with hierarchical clustering by Euclidean distance within each cluster. C) GO analysis of the clusters from the top 2000 highly variable genes from B. The size of the circles indicates the gene ratio (number of genes from the input list annotated to the GO term divided by the total number of genes in the input list), and the color represents the significance of the adjusted p-value. Gene clusters are color-coded on the y-axis and GO processes are shown on the x-axis.

Of the 2000 most variably expressed genes among the HL-60 samples, only 215 had increased expression specific to hypomethylated colonies (Fig. 4B, cluster 3; Supplementary Table 9). Many of these genes (42.3%) were upregulated after 72h of treatment with either DAC or AZA in bulk HL-60 RNA-seq data, and several were associated with activation of inflammatory responses within the sPLS model (e.g., S100A8 and S100A9; Supplementary Table 9; Fig. 2B, cluster 1). In contrast, the genes in cluster 5 were upregulated following HMA exposure in both hypomethylated and highly methylated colonies (Fig. 4B), and only 3.5% of these genes were induced by HMA treatment in bulk RNA-seq data (Supplementary Table 9). Thus, some transcriptional changes that occur immediately after HMA treatment are maintained in hypomethylated colonies, whereas other genes are upregulated during colony formation and are independent of HMA-induced global hypomethylation. GO ORA revealed an enrichment of anti-microbial and immune-related processes among both hypomethylation-dependent and -independent gene sets, whereas ‘small molecule biosynthetic process’ and ‘cholesterol biosynthetic process’ were over-represented among the hypomethylation-independent cluster 5 genes (Fig. 4C; Supplementary Table 10). HL-60 colonies that retained DNA methylation following DAC treatment also had particularly high expression of cholesterol-related genes, including many enzymes required for de novo cholesterol biosynthesis downstream of mevalonate (34) (Supplementary Fig. S6-7; Tables S11, S12).

We next performed a focused analysis of genes annotated to ‘cholesterol biosynthetic process’ (GO:0006695) across all cell lines. Like HL-60 colonies, MOLM-13 and MV-4-11 samples displayed increased expression of many of these genes after HMA treatment (Supplementary Fig. S8).

Furthermore, a statistical analysis revealed that 5 genes (SREBF1, SCAP, PMVK, MVD and LSS) were significantly increased (Pairwise Wilcoxon test with Benjamini-Hochberg correction, FDR = < 0.05) by both DAC and AZA in colonies derived from all 3 cell lines (Fig. 5A). These genes form part of a regulatory feedback loop that maintains cholesterol homeostasis by up-regulating biosynthesis in response to low cellular cholesterol (35) (Supplementary Fig. S6). Since cholesterol regulation is altered in both methylation-retaining and hypomethylated colonies, targeting this metabolic pathway could overcome the initial heterogeneity induced by HMA treatment (Fig. 1-3).

**Figure 5.**
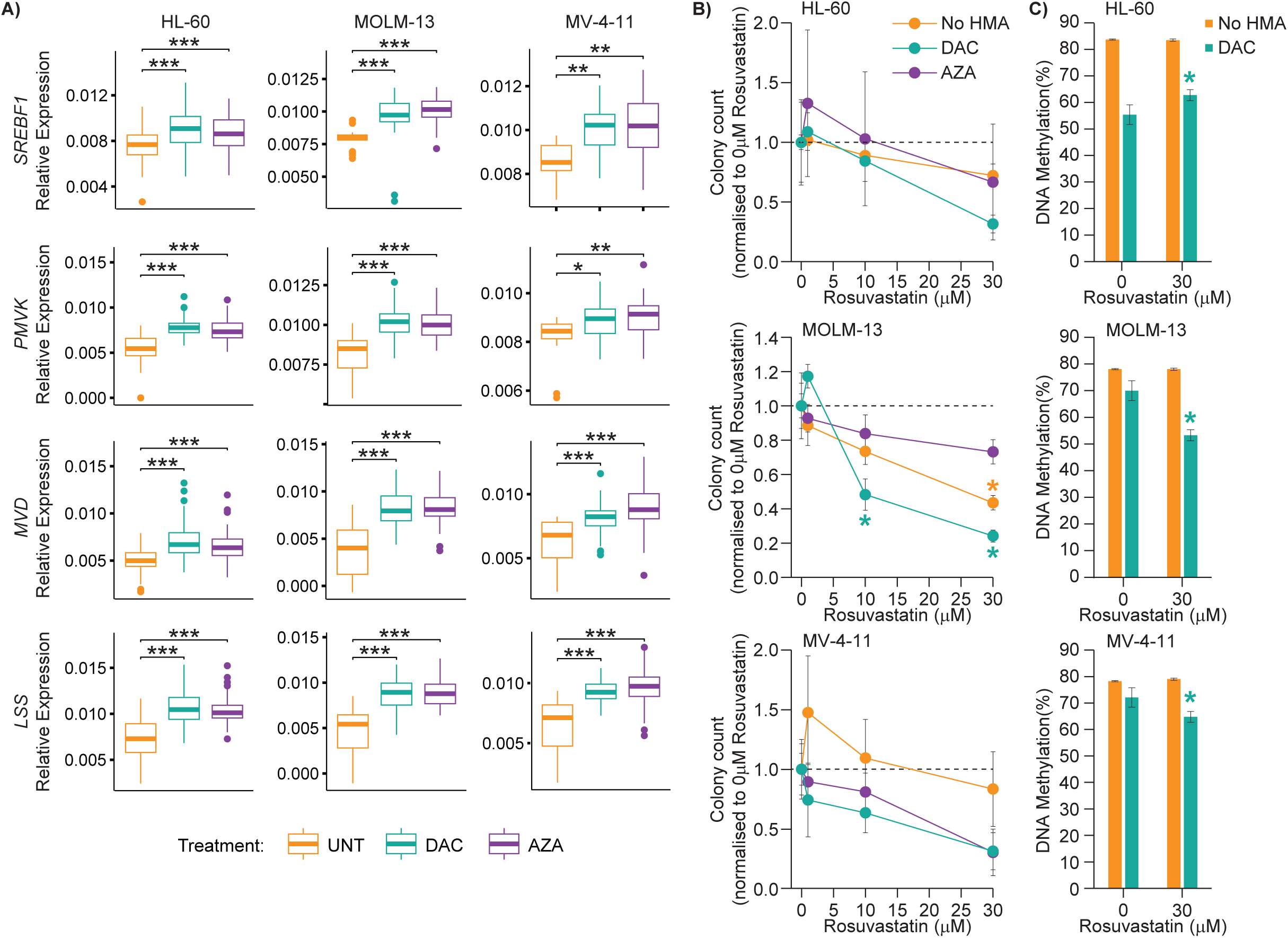
Altered cholesterol regulation in cells surviving HMA therapy can be targeted to enhance treatment efficacy. A) Box and whisker plots of relative expression for SREBF1, PMVK, MVD and LSS in HL-60 (left), MOLM-13 (middle) and MV-4-11 (right) colony data from Figure 5. Pairwise Wilcoxon test between treatment groups was performed with UNT samples as the reference group, for genes present within ‘cholesterol biosynthetic process’ (GO:0006695). Boxes depict the interquartile range (IQR) with median. Whiskers extend to the highest and lowest data points within 1.5 x IQR of the first and third quartile. Benjamini-Hochberg false discover rate (FDR) correction was performed on all p-values, per cell line: FDR < 0.05*, FDR < 0.01**, FDR < 0.001***. B) Colony counts for HL-60 (top), MOLM-13 (middle) and MV-4-11 (bottom) cell lines obtained following HMA and rosuvastatin co-treatments. Data from No HMA (orange), DAC (cyan) and AZA (purple) groups are normalised to the corresponding 0 μM rosuvastatin control. Means ± SEM for n = 3 experiments. Significance determined by two-way ordinary ANOVA with Dunnett’s multiple comparisons test, p < 0.05* vs. corresponding 0 µM rosuvastatin control. C) DNA methylation of colonies formed following DAC and rosuvastatin co-treatments. Means ± SEM for n = 3 experiments. Significance determined by one-way ANOVA with Tukey’s multiple comparisons test, p < 0.05* vs. corresponding 0 µM rosuvastatin control.

### Inhibition of cholesterol pathways following HMA exposure

To further investigate the relationship between cholesterol regulation and self-renewal capacity following HMA treatment, we performed colony-forming assays in the presence of rosuvastatin. Rosuvastatin is a potent inhibitor of the rate-limiting enzyme of the cholesterol biosynthetic pathway, HMG-CoA reductase (HMGCR), and is frequently prescribed to reduce the risk of cardiovascular diseases associated with hypercholesterolaemia (36). AZA and rosuvastatin co-treatment had varied effects in AML cell lines (Fig. 5B, C), with synergistic effects on colony-formation observed in MV-4-11 cells (Supplementary Fig. S5C). Meanwhile, DAC and rosuvastatin had synergistic effects on colony formation in all cell lines (Supplementary Fig. S5C), with significantly fewer colonies formed upon addition of 10 or 30 µM rosuvastatin in MOLM-13 cells (Fig. 5B). Colonies also displayed significantly reduced size (Supplementary Fig. S5B) and DNA methylation levels (Fig. 5C) following DAC and rosuvastatin treatment of MOLM-13 and MV-4-11 cell lines, where strong selection for highly methylated cells was previously noted (Fig. 3B). This suggests that inhibition of cholesterol biosynthesis is effective even against methylation-retaining cells that maintain self-renewal capacity following HMA treatment.

The efficacy and potential clinical utility of DAC and rosuvastatin co-treatment was then tested in immunocompromised mice engrafted with luciferase-tagged MOLM-13 cells. In a dose optimization experiment, combining DAC and rosuvastatin led to a significant increase in median survival for all doses tested (Fig. 6A; Supplementary Table 13). The 1 mg/kg/day dose of rosuvastatin was chosen for further validation in this model, as it is a low dose commonly prescribed for preventing cardiovascular disease (equivalent to approximately 5mg/day oral dose in humans). Co-treatment with DAC and rosuvastatin once again led to a significant increase in median survival compared to DAC treatment alone (24 vs. 21 days) in this aggressive in vivo AML model (Fig. 6B; Supplementary Table 14). In contrast, rosuvastatin alone significantly reduced the median survival time of vehicle-treated mice (18 vs. 19 days; Fig. 6B; Supplementary Table 14). Finally, we assessed AZA and DAC in combination with rosuvastatin in a patient-derived xenograft model of AML (AML-16). Rosuvastatin (1mg/kg/day) co-treatment significantly extended the median survival of AZA-treated mice by 8.5 days (Fig. 6C; Supplementary Table 15) and decreased leukemia burden (Fig. 6D). Meanwhile, combining rosuvastatin with DAC (0.2mg/kg/day) had no significant effect on survival or leukemia burden in this model (Supplementary Fig. S9). These in vivo results provide evidence that HMAs combined with statins have the potential to decrease leukemia burden and improve AML survival, with further research required to optimize drug selection and dosing in patient cohorts.

**Figure 6.**
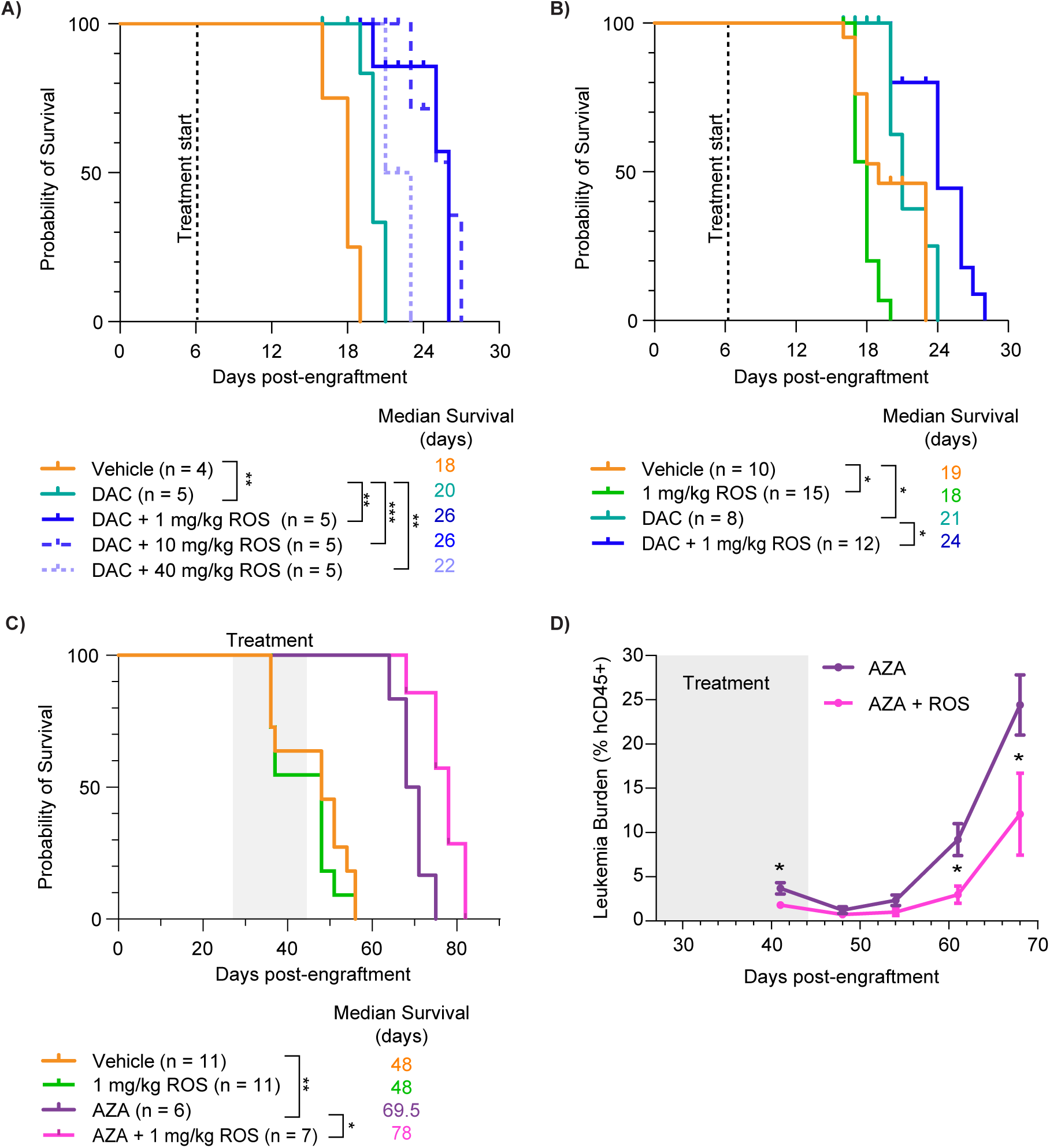
HMA and statin co-treatment increases survival in vivo. A) Rosuvastatin dose optimisation experiment showing median survival of mice engrafted with MOLM-13-luc cells following treatment with DAC (0.2mg/kg/day) +/- rosuvastatin (1, 10, 40mg/kg/day) on a treatment schedule of ‘5 days on, 2 days off’ for 3 cycles. B) Validation of survival benefit when DAC (0.2mg/kg/day) is combined with rosuvastatin (1mg/kg/day) in mice engrafted with MOLM-13-luc AML cells. C) AZA and rosuvastatin co-treatment significantly extended median survival of NSG mice engrafted with a patient-derived xenograft (AML-16 model). Mice were treated with AZA (1mg/kg/day) +/- rosuvastatin (1mg/kg/day) for 5 days on, 2 days off in cycle 1, followed by twice per week for an additional 2 cycles. D) Leukemia burden (% human CD45^+^ cells) post-engraftment and following the cessation of treatment. Mann-Whitney test (unpaired, non-parametric, two-tailed t-test) was used to test for statistical significance. Survival analyses (A-C) were performed using Kaplan-Meier analysis followed by the Log-rank (Mantel-Cox) test and an adjusted p-value of < 0.05 was considered statistically significant. p < 0.05*, p < 0.005**, p < 0.0001***.

## Discussion

A thorough understanding of the ways in which AML cells survive HMA therapy and initiate recurrence is essential to overcome the clinical challenge of treatment-resistant relapse. Our unique study has revealed HMA-induced heterogeneity and methylation-retaining cells that evade the epigenetic effects of treatment. We also identify metabolic pathways associated with the self-renewal of HMA-treated AML cells, which can be targeted to improve HMA effects.

Our single-cell multiomic analyses revealed global DNA methylation heterogeneity induced by treatment and transcriptional responses linked to epigenetic changes. We observed activation of inflammatory response and cell death pathways in only a minor subset of hypomethylated cells (Fig. 2B), consistent with our previous report of heterogeneous TE expression following DAC treatment (24) and scRNA-seq data from a colorectal cancer cell line (37). These observations suggest that additional epigenetic modifications, transcription factors, or other mechanisms can suppress transcriptional responses in hypomethylated cells (e.g., H3K9me3 (38, 39)). Alternatively, loss of methylation at specific loci may be required for HMAs to exert their effects. Our observations are consistent with the lack of correlation between HMA-induced global hypomethylation and clinical response (40, 41) and support the use of locus-specific methylation changes to build a predictor of patient response (42).

In contrast to our single-cell analyses, Li et al. reported reduced epigenetic and transcriptional variance in AML cells collected after 12 weeks of AZA treatment in a transgenic mouse model (43). We suggest that HMAs initially increase epigenetic and transcriptional diversity, allowing some cells to gain a relative growth or survival advantage. Subsequent expansion of these clones would lead to reduced heterogeneity, as reported by Li et al. (43). Consistently, we found that some AML cells retained high levels of DNA methylation during HMA treatment (Fig. 1A) and had a relative growth advantage following drug withdrawal (Fig. 3). Methylation-retaining cells tended to divide less frequently during treatment (Fig. 1B) but had higher self-renewal and proliferative capacity than hypomethylated cells after DAC treatment (Fig. 3). Interestingly, this selection for methylation-retaining cells was less pronounced following AZA treatment, reflecting previous reports of distinct effects of these HMAs (44, 45). Since AZA is incorporated into both DNA and RNA (46), whereas DAC is restricted to DNA, we speculate that RNA-mediated toxicities (such as translational inhibition (14, 15)) could prevent the growth of highly methylated cells following AZA treatment. Taken together, our results show that heterogeneous responses to HMA treatment can influence AML cell self-renewal with potential implications for relapse.

We observed upregulated expression of cholesterol biosynthesis genes in colonies formed after HMA treatment (Fig. 5; Supplementary Fig. S6, 8). These changes were observed in both hypomethylated and highly methylated colonies, suggesting that HMA exposure has persistent transcriptional effects that are independent of global methylation loss. Rapidly proliferating AML cells are known to have high cholesterol demands (47), with cholesterol and related metabolites required for cell membrane synthesis, lipid rafts, cell cycle progression and activation of oncogenic signaling molecules (via prenylation of RAS, Rho/Rac) (Supplementary Fig. S7) (34, 48). In colony-forming assays rosuvastatin synergistically enhanced the effects of DAC (Fig. 5B; Supplementary Fig. S5C), suggesting that rapid consumption of cholesterol supports AML cell self-renewal after HMA treatment (35). Since this metabolic adaptation occurs even in highly methylated cells, statin co-treatment could have broad impact to inhibit AML cell growth despite initial heterogeneity in HMA responses.

Altered cholesterol regulation is an established feature of AML with relevance to prognosis and therapy (47, 49, 50). Recently, cholesterol metabolism was also linked to DAC resistance in AML cell lines, with statin co-treatment showing synergistic inhibition of in vitro AML cell growth (51). Others have shown that inhibition of cholesterol biosynthesis can sensitize AML cells to radiation and chemotherapy (52, 53). In vivo, we observed significantly improved survival of leukemia-bearing mice treated with HMA and rosuvastatin (Fig. 6) suggesting that co-treatment may increase the duration of response in some AML and MDS patients. Encouragingly, a recent retrospective analysis of MDS patients (including some HMA-treated high-risk cases) reported improved survival and reduced progression to AML in patients who commenced statin treatments three months before or after MDS diagnosis (54). Current clinical trials are testing the safety of a statin (pitavastatin) in combination with AZA and venetoclax in AML patients (55). This therapeutic avenue is of particular interest since statins are commonly prescribed, well-tolerated, oral medications, which could be rapidly repositioned for use in AML and MDS patients receiving HMA therapy. Our results support further investigation of the long-term benefits of statin and HMA co-treatments in different AML subtypes.

## Methods

### Cell lines and culture

AML cell lines (Supplementary Table 1), HL-60 (ATCC #CCL-240), MOLM-13 (DSMZ #ACC-554), and MV-4-11 (ATCC #CRL-9591), were maintained in tissue culture flasks (Greiner Bio-One) at 37°C and 5% CO_2_, and sub-cultured at 500,000 cells/mL every 2-3 days with fresh medium. HL-60 cells were maintained in Iscovellls Modified Dulbeccollls medium (IMDM; Sigma-Aldrich) supplemented with 4mM GlutaMAX (Thermo Fisher Scientific) and 10% Fetal Bovine Serum (FBS; Sigma-Aldrich). MOLM-13 and MV-4-11 cells were maintained in Roswell Park Memorial Institute 1640 medium (RPMI; Sigma-Aldrich) supplemented with 2mM GlutaMAX and 10% FBS.

### CellTrace staining

AML cells (2-2.5×10^6^ cells/mL) were stained with 1µM (MOLM-13 and MV-4-11) or 3µM (HL-60) CellTrace Far Red (Thermo Fisher Scientific), according to the manufacturer’s instructions. MOLM-13 and MV-4-11 cells with uniformly high CellTrace fluorescence were purified by fluorescence-activated cell sorting (FACS) prior to treatment with hypomethylating agents (HMAs), whereas all cells were used for HL-60 treatments.

### HMA treatments

HMA treatments were performed in suspension culture, with drugs added every 24h for 72h total. All cell lines were treated with 100nM decitabine (DAC; Selleckchem #S1200), HL-60 cells were treated with 2000nM azacytidine (AZA; Selleckchem # S1782), while MOLM-13 and MV-4-11 cells were treated with 500nM AZA. Untreated cells (UNT) received 0.1% DMSO (vehicle control).

### Fluorescence activated cell sorting (FACS)

HMA-treated cells were stained with propidium iodide (PI, 1.5µg/mL) and prepared for FACS. Viable (PI^-^) single cells were sorted into 2.5µL of RLT PLUS buffer (Qiagen) containing 2.5U SUPERas-In (Thermo Fisher Scientific) in 96-well plates using indexed sorting on a FACS Aria II (BD Biosciences). Plates were sealed and briefly centrifuged before storage at −80°C for sequencing analysis.

### Colony-forming assays

HMA-treated cells were seeded in MethoCult Optimum (H4034; STEMCELL Technologies Inc.) at 500 cells/well in 6-well plates with rosuvastatin (Selleckchem # S2169) added to the MethoCult at various doses (0, 1, 10, 30µM). Cells were cultured at 37°C and 5% CO_2_ for 14 days, and colonies were imaged using Cytation3 (Biotek). Colony counts and sizes were analyzed using ImageJ software. Individual colonies were manually plucked using a 20µL pipette tip into 100µL of media, centrifuged at 200xg for 5 min, and then resuspended in 20µL of RLT PLUS buffer before storage at −80°C. Alternatively, all colonies in each well were collected by resuspending the MethoCult Optimum media (and colonies) in 3mL of standard culture media (IMDM or RPMI), centrifuging at 200xg for 5 min, and resuspending the cell pellet in 20-50µL of RLT PLUS buffer before storage at −80°C.

### Library preparation and sequencing

#### scNMT-seq library preparation and sequencing

For scNMT-seq, matched scNOMe-seq and scRNA-seq libraries were prepared from sorted HL-60 single cells as previously described (23). Minor modifications to the published protocol were as follows: 1) Amplified cDNA was purified using a 0.6:1 volumetric ratio of AMPure XP Beads (Beckman Coulter) and eluted in 15µL water. 2) During the fifth repeat of first-strand synthesis reaction, the samples were held at 37°C for 90min. 3) The oligo used in the second-strand synthesis reaction (5’-CAGACGTGTGCTCTTCCGATCTNNNNNN-3’) was modified to be compatible with the NEBNext oligos (multiplex oligos for Illumina, dual index sets, New England Biolabs), which were used to amplify scNOMe-seq libraries. 4) A 0.65:1 volumetric ratio of AMPure XP Beads was used to purify products of both the first- and second-strand synthesis reactions, as well as the amplified scNOMe-seq libraries.

For scNOMe-seq libraries, paired-end 150bp sequencing was performed using the NovaSeq (Illumina) platform. For scRNA-seq libraries, paired-end 75bp sequencing was performed on the NovaSeq or NextSeq (Illumina) platform.

#### scTEM-seq library preparation and sequencing

For scTEM-seq analysis of global DNA methylation levels in single MOLM-13 and MV-4-11 cells, library preparation was performed as described (24). Paired-end 150bp sequencing was performed on the MiSeq (Illumina) platform.

### Colony TEM-seq and RNA-seq library preparation and sequencing

Single-colony TEM-seq (Fig. 3B, C) and parallel RNA-seq analyses were performed as described (24, 25) with minor modifications. Lysates from single colonies (HL-60: 7.5µL; MOLM-13 and MV-4-11: 2.5µL) were used to separate genomic DNA and mRNA. During single-colony TEM-seq library preparation, the number of SINE Alu amplification cycles was reduced to 29. For RNA-seq analysis, 15 cycles of cDNA amplification were performed. TEM-seq analysis of pooled colonies (Fig. 3D, 5E) was performed as described (25) using 5-10µL of cell lysate as input for bisulfite conversion and 29 cycles for SINE Alu amplification.

TEM-seq libraries were sequenced using 150bp paired-end sequencing on the MiSeq platform. For RNA-seq libraries, paired-end 75bp sequencing was performed on the NextSeq or NovaSeq platform.

### Data processing

Sequencing data were processed and aligned as described in the Supplementary Information.

### scNMT-seq data analysis

Quality control, data normalization and batch correction were performed as described in the Supplementary Information. All analyses were performed in R v4.2.1, unless otherwise stated.

### Pairwise distance analysis of DNA methylation heterogeneity

To assess DNA methylation heterogeneity per treatment group and genomic context (Fig. 1D), pairwise CpG methylation distance analysis was performed. The mean absolute methylation difference was computed for each cell pair (A, B) as the mean of the absolute difference in methylation rate at each common cytosine position in the relevant genomic annotation. To make the comparison of methylation patterns meaningful, only cytosine loci with data from both cells in the pair were used. The mean absolute methylation differences were grouped according to the treatment combination of the cell pairs. The global summaries shown in Fig. 1D correspond to groups in which both cells in the pair had the same treatment. Higher values indicate a more heterogeneous methylation pattern when cells in the same treatment group were compared vis-à-vis.

### Cell-wise correlation analysis

To assess the relationships between DNA methylation, DNA accessibility, and gene expression within individual cells (Fig. 1E), Pearson correlations were computed using HL-60 scNMT-seq data. For this analysis, RNA-seq data was normalized and log transformed per batch using ‘scuttle::logNormCounts()’ (v1.6.2) (26) without batch correction or prior count filtering. DNA methylation was correlated to DNA accessibility at matched loci, based on genomic co-ordinates. For correlations with gene expression, methylation and accessibility measurements at promoters, introns, and exons were matched to the corresponding transcript. For CpG islands, H3K27ac sites and H3K4me3 sites, methylation and accessibility measurements were matched to all transcripts within 10kb. For intergenic regions and 3kb genomic windows, methylation and accessibility measurements were matched to every transcript within 1bp to assess the expression of immediately adjacent genes. For each cell, Pearson’s correlation estimates were then computed using all matched values and the ‘cor.test()’ function in R.

### Integrative sparse partial least squares (sPLS) analysis

Mixomics (27) (v6.20.0) was used to perform a multivariate integrative analysis of the HL-60 scNMT-seq data (Fig. 2A-D). Feature selection was performed to identify variably expressed transcripts that were highly correlated with changes in DNA Methylation and accessibility after HMA treatment. We performed an unsupervised sparse Partial Least Squares (sPLS) analysis using the function ‘mixOmics::mint.block.spls()’ which combines a multivariate integrative (MINT) method and a multiblock sPLS integrative analysis. MINT (28) accounts for multiple batches (Supplementary Table 2) measured on the same variables, while the multiblock sPLS seeks for correlated patterns between DNA methylation and DNA accessibility rates that are split into multiple genomic regions (‘blocks’) and explain (correlated to) the predictor (transcriptome).

To focus on transcriptomic and epigenetic changes resulting from HMA treatment, only treated cells (AZA and DAC) were included in the sPLS model. The genomic regions included in this analysis for both DNA methylation and DNA accessibility were CpG islands, promoters, H3K27ac sites, H3K4me3 sites and 3kb windows. The rates from these genomic regions were filtered to retain only those detected in greater than 10% of cells. The sPLS model was implemented assessing two components, selecting 50 features per component and per block (genomic region) in the DNA methylation and DNA accessibility datasets, and 100 genes per component in the transcriptome dataset. Further details are provided on GitHub.

Heatmap visualization and identification of cell and expression clusters from sPLS-selected features was performed using the ComplexHeatmap package (v2.12.1). sPLS selected features for components 1 and 2 were extracted using the function ‘mixOmics::selectVar()’. Heatmap visualization was performed on sPLS-selected features using transcriptomic (converted to z-score) and epigenetic rates (mean of features in genomic regions and converted to z-score) that were entered into the model and included both treated and untreated cells. K-means clustering was performed on sPLS-selected transcriptomic features, first on Gene Expression (row_km=3), followed by Cell Group (column_km=4).

sPLS sample projections (Fig. 2A, C, and D) were plotted using ggplot2 (29) (v3.3.6) by extracting the sPLS components 1 and 2 for a given block (RNA or epigenetic genomic region) and overlaid with relevant information, that is, the cell group identified from k-means clustering and treated cell type or average DNA methylation.

Gene Ontology (GO) Over Representation Analysis (ORA) was calculated using clusterProfiler (30) (v4.4.4) for ‘biological process’ and displayed using enrichplot (v1.16.1) (Fig. 2E, F). Gene Expression k-means clusters (Fig. 2B) and sPLS selected features per component (1–2) were assessed by ‘enrichGO(p.adj=0.05, p.adj.method = “fdr”, q.val.threshold = 0.4)’ and the list of genes from the batch corrected transcriptome dataset (entered into the sPLS model) as the background. Results were displayed as treeplots using default settings for pairwise ‘termsim()’ and ‘treeplot(nCluster=5, showCategory = 10)’.

The correlation of sPLS features (Supplementary Fig. S3) was calculated as a similarity matrix using ‘mixOmics::circosPlot()’ on the sPLS model. The results were displayed using ComplexHeatmap, showing DNA methylation and DNA accessibility features related to transcript features split by the previously identified Gene Expression k-means clusters.

### Locus-specific correlation analysis

To compare gene expression to adjacent epigenetic features, locus-specific correlations were performed using HL-60 scNMT-seq data (Fig. 2H, I). DNA methylation and DNA accessibility measurements were paired to genes based on annotation (promoters) or by strand-aware position within 10kbp of the gene transcription start site (CpG islands, H3K27ac sites, H3K4me3 sites and 3kb windows). For paired sites, Pearson’s correlations were computed between CpG or GpC methylation rate and log gene expression values. All cells with data (i.e., the UNT, DAC, and AZA groups) were combined in these correlations, and a minimum of 22 cells with paired data (i.e., both gene expression and DNA methylation/accessibility measurements) were required for the correlation to be performed.

### Colony sequencing analysis

Quality control, data normalization and batch correction were performed as described in the Supplementary Information. All analyses were performed in R v4.2.1.

### Highly variable gene analysis

For Figure 4, highly variable genes (HVGs) were identified from colony RNA-seq data and principal component analysis (PCA) was performed using ‘scater::calculatePCA(ntop = 2,000)’. K-means clusters of HVGs were determined using the R stats package (v4.2.1) with ‘kmeans(centers = 8, iter.max = kmeans.iter, nstart = 50)’. Heatmapping of HVGs and k-means cluster was performed using ‘ComplexHeatmap::pheatmap()’ with z-scored values and Euclidean distance hierarchical clustering within row clusters (k-means groups) and columns (samples) ordered by treatment and descending average global methylation level.

GO ORA of k-means clusters was compared using clusterProfiler for ‘biological process’ by ‘compareCluster(pAdjustMethod = “fdr”, p.adj.threshold = 0.05, qvalueCutoff=0.4)’ and the full list of genes from the batch corrected dataset (for each cell type) as the background list. Plots were created using ‘clusterProfiler::dotplot(showCategory = 3) + coord_flip()’.

### Analysis of cholesterol biosynthesis gene expression

For Figure 5A, genes from the cholesterol biosynthesis pathway (GO:0006695) were analyzed from batch corrected expression data for each cell line (HL-60, MOLM-13 and MV-4-11). After batch correction, 40 genes out of 58 genes (from GO:0006695) were in common across all three cell lines datasets. In total, for each cell line there were 41 (HL-60), 41 (MOLM-13) and 43 (MV-4-11) genes analyzed from GO:0006695. The R package rstatix (v0.7.2) was used to perform pairwise Wilcoxon test between treatment groups with UNT samples as the reference group for each gene. Benjamini-Hochberg false discovery rate (FDR) correction was performed on all p-values, per cell line. Boxplots were generated using ggpubr (v0.6.0) displaying the batch corrected relative expression with ‘pointrange’ error.

### AML xenograft models

Five-week-old female NOD.Cg-Prkdc scid Il2rg tm1Wjl /SzJ (NSG) mice were obtained from the Australian Bioresources (ABR, Moss Vale, NSW, Australia) and acclimatized for one week prior to any experimental procedure.

MOLM-13 cells transduced with firefly luciferase (MOLM-13-luc; 5×10^5^ cells suspended in 100 μL of PBS) were injected into the lateral tail vein of NSG mice. Leukemia burden was assessed by bioluminescence imaging (BLI) twice a week using an IVIS Spectrum in vivo imaging system (PerkinElmer, Waltham, MA, USA), following intraperitoneal injections of 150 mg/kg D-luciferin (Promega, Alexandria, NSW, Australia) under anesthesia with isoflurane. Treatments commenced on day 6 post-engraftment when a positive luminescence signal was detected. Mice (n = 5-15/group) were treated by intraperitoneal injection of either vehicle (2% DMSO, 30% PEG300 in water), DAC (0.2 mg/Kg in saline), rosuvastatin (1 mg/kg), or combination rosuvastatin (1 mg/kg, 10 mg/kg, or 40 mg/kg in 30% PEG300 in water) and DAC (0.2 mg/Kg) once a day (5 days on, 2 days off) for up to 3 weeks. The animals were monitored until they reached the ethical endpoint.

For patient derived xenograft (PDX) experiments, NSG mice were inoculated with 2×10^6^ AML-16 cells (FLT3-ITD, NPM1, IDH2, and WT1 mutant) (31) in 100µL of PBS, by injection into the lateral tail vein. Leukemia burden (% hCD45+ cells in peripheral blood) was assessed 1-2 times weekly via flow cytometry (FACSCanto II). Treatment commenced once an average leukemia burden of 1% CD45^+^ cells was achieved (∼3.5 weeks). Mice (n= 6-11 per group) received vehicle (2% DMSO, 30% PEG300 in water), DAC (0.2 mg/Kg), AZA (1mg/Kg), rosuvastatin (1 mg/kg), DAC + rosuvastatin, or AZA + rosuvastatin, via intraperitoneal injections once a day (5 days on, 2 days off) for 1 week (cycle 1), followed by twice per week for an additional 2 weeks (2 cycles). Mice were monitored until the ethical endpoint (>25% huCD45^+^ cells).

### Survival Analysis

Survival analyses were performed using Kaplan-Meier analysis followed by the log-rank (Mantel-Cox) test. All statistical analyses were performed using GraphPad Prism v. 9.0 (GraphPad Software, La Jolla, CA, USA). Differences were considered statistically significant at an adjusted p-value < 0.05.

## Data Availability

Sequencing data generated in this study has been deposited in the GEO database (GSE255339). Relevant code for this manuscript is available on GitHub via: www.github.com/canepi/HMA_heterogeneity

## Acknowledgements

Nicole Cole (University of Newcastle) provided technical support for fluorescence activated cell sorting. Al J. Abadi (Melbourne Integrative Genomics, University of Melbourne) assisted with the implementation of sPLS analysis. The authors thank Professor Tri Phan (Garvan Institute) and Professor Xu Dong Zhang (University of Newcastle) for critically reviewing this manuscript. H.J.L., C.R. and M.D.D. received funding from the National Health and Medical Research Council of Australia (NHMRC) (GNT1143614, GNT1180782, GNT1173892, and GNT2016283). K.A.L.C. and H.J.L. received funding from the Australian Research Council (ARC) (DP200102903). H.J.L. and H.C.M. received funding from the Cancer Institute NSW (2018/ECF001 and ECF1299, respectively). D.R.B received funding from Cure Cancer Australia Foundation (CCAF2023-Bond) and NSW Health Pathology North. N.A.B. received funding from the McGuigan family through the Hunter Medical Research Institute (HMRI778). The contents of the published material are solely the responsibility of the research institutions involved or individual authors, and do not reflect the views of funding agencies.

## Author Contributions

H.J.L. conceived and oversaw the project. D.R.B., A.K.E., C.R., K.A.L.C., and H.J.L. acquired funding. D.R.B., K.U., K.V.H., B.H., C.L.O., E.A.R., S.H., L.S.R., H.M., and H.J.L. performed experiments. D.R.B., S.M.B., K.U., K.V.H., B.H., C.L.O., L.S.R., C.R., and H.J.L. performed data analysis. S.M.B. and C.R. developed analytical approaches. S.M.B. managed sequencing data and analysis code. D.R.B., N.A.B., A.K.E, N.M.V., K.A.L.C., and H.J.L. supervised work. A.M., M.D.D. and C.E.d.B. provided critical resources. D.R.B., S.M.B., K.U., B.H., L.S.R., C.R., and H.J.L. prepared figures and wrote the manuscript. All authors have read and approved the manuscript.

## Ethics Declarations

All experimental procedures using animals were reviewed, approved, and performed according to the Animal Care and Ethics Committee of the University of Newcastle (approval number: A-2023-303), and with consideration of the ARRIVE guidelines (Supplementary Information).

## Notes

### Competing Interest Statement

A.K.E. declares the following competing interests: advisory board and honoraria from AbbVie, Astellas, Gilead, Servier, Jazz, Otsuka/Astex, and RACE Oncology; speaker fees from AbbVie, Otsuka/Astex, Astellas and Jazz; Research Funding from RACE oncology and Otsuka/Astex. N.A.B. receives research funding from Merck KGaA and Bristol Myers Squibb.

### Summary of Updates

Figure 5 revised. Inclusion of new data from in vivo cancer models in Figure 6.

