## Supplementary Material for "Hypomethylating agents induce epigenetic and transcriptional heterogeneity with implications for AML cell self-renewal"

### Supplementary Information

#### Supplementary Methods

#### Supplementary References

#### Supplementary Figures S1 to S9

#### Supplementary Tables S1, S7, S13, S14, S15

#### Supplementary Methods

##### Bulk RNA sequencing

HL-60 cells were treated every 24h for 72h with HMAs (100nM DAC, 2000nM AZA). Cells (1x10^6^) were stored in 1x DNA/RNA Shield (Zymo Research) and RNA was extracted using the Quick DNA/RNA MagBind kit (Zymo Research), according to manufacturer’s instructions. RNA was quantified on a Nanodrop, and integrity was assessed using a TapeStation RNA ScreenTape. RNA-seq libraries (700ng RNA input) were prepared using the TruSeq Stranded mRNA kit (Illumina), according to the manufacturer’s instructions, and sequenced on the NovaSeq 6000 system (SP flow cell; 150 cycles). Sequencing data was analysed using Seqmonk (QC and DESeq2).

##### Post-bisulphite adaptor tagging (PBAT) library preparation, processing and analysis

DNA methylation analysis presented in Supplementary Fig. S1 was performed using a post-bisulphite adaptor tagging (PBAT) sequencing approach as previously described (1).

##### Bulk TEM-seq library preparation and sequencing

For analysis in Supplementary Fig. S2, CellTrace-low (bottom 30-35%), -high (top 15-20%) and -medium (middle 40-50%) populations were purified by FACS and resuspended in 30µL RLT PLUS buffer (Qiagen) containing 30U SUPERas-In (Thermo Fisher Scientific). Lysates were stored at -80 °C for TEM-seq analysis.

TEM-seq analysis of CellTrace-low, -medium and -high populations was performed as described (1, 2) using 5-10µL of lysate as input for bisulfite conversion and 25 cycles for SINE Alu amplification.

##### scNOMe-seq data processing

Single-cell bisulfite libraries were processed using Bismark (3) (v0.22.3) as described (4). Following demultiplexing, samples with multiple sequencing lanes per read were concatenated into single R1 and R2 fastq files per sample. Reads were trimmed using Trim Galore (5) (v0.6.6) with cutadapt (v1.18) to remove automatically detected illumina adaptors and the first 9 reads ‘--Clip_R1 9’ in single end mode. Reads were then mapped by single-end non-directional alignment to Bowtie2 indexed human genome GRCh38. Sample reads were deduplicated in single-end mode and concatenated into a single file per sample by ‘deduplicate_bismark --multiple --single’. Methylation extraction and coverage file generation was performed separately on samples with ‘bismark_methylation_extractor --single-end --bedGraph --gzip --cx’, followed by ‘coverage2cytosine --gzip --nome --seq’. For analyses comparing HL-60 scNOMe-seq data and colony TEM-seq data (Fig. 3B), scNOMe-seq data was filtered for CpG sites overlapping SINE Alu sites.

##### (sc)TEM-seq data processing

Bisulfite sequencing data processing and extraction of global methylation estimates for both colonies and single-cell samples were performed as previously described (1, 2) and scripts available via <https://github.com/canepi/scTEM-seq>. In scTEM-seq data from MOLM-13 and MV-4-11 cells, samples were excluded if fewer than 500 unique SINE Alu annotations were represented.

##### (sc)RNA-seq data processing

Following demultiplexing, multiple sequencing lanes per read were concatenated into single R1 and R2 fastq files, per sample. Raw reads were trimmed in pair-end mode using Trim Galore (v0.6.5) and Cutadapt (v2.10), retaining unpaired reads. Hisat2 (6) (v2.1.0) was used to map and align trimmed and unpaired reads using default parameters (--phred33) to the human reference genome build (GRCh38), and samtools (v1.10) was used to sort by coordinate and generate bam files.

TEtranscripts (7) (v2.2.1) was used to obtain raw gene and transposable element counts from unique and ambiguously aligned reads (bam files) using TEcount with the flags '--sortByPos --mode multi’. The GTF files used were generated as follows: 1) TEs (<http://hgdownload.soe.ucsc.edu/goldenPath/hg38/database/rmsk.txt.gz>; USCS repeating element Repeat Masker; hg38; 2021-09-03) and 2) genes (<ftp://ftp.ebi.ac.uk/pub/databases/gencode/Gencode_human/release_30/gencode.v30.annotation.gtf.gz>; gencode Release 30; GRCh38.p12; 2019-04-08). To ensure the GTF files worked with
TEtranscripts the gene GTF (from gencode) was updated by converting ‘M’ to ‘MT’ for mitochondrial genes in the chromosome column and removal of ‘Chr’ preceding each chromosome number. The TE GTF was generated from the repeat mask file using the conversion tool from the TEtranscripts developers (<http://labshare.cshl.edu/shares/mhammelllab/www-data/TEtranscripts/TE_GTF/makeTEgtf.pl.gz>) ‘perl makeTEgtf.pl -c 6 -s 7 -e 8 -o 10 -t 11 -n hg38_rmsk -f 13 -C 12 -S 2 rmsk.txt >> hg38_rmsk_TE.gtf’ and removing ‘chr’ from chromosome names ‘sed 's/chr//' hg38_rmsk_TE.gtf > hg38_rmsk_TE_NoChr.gtf’.

##### scNMT-seq quality control

For scNMT-seq data, cells were required to pass both scNOMe-seq and scRNA-seq quality control (QC). Cells that had less than 500,000 CpG sites covered, less than 5,000,000 GpC sites covered, greater than 15% CHH methylation rate, or less than 2% GpC methylation failed scNOMe-seq QC. For scRNA-seq, QC was performed using bam files from hisat2 and the SeqMonk (v1.47.1) ‘RNA-seq QC Plot’. Cells that had less than 70% reads in exons or less than 15% genes measured failed scRNA-seq QC. In total, 222 scNMT-seq samples passed QC (Dataset S1, Table S2).

##### scNOMe-seq normalisation and batch correction

scNOMe-seq libraries provide information on both DNA methylation (CpG sites) and DNA accessibility (GpC sites). For both CpG (methylation) and GpC (accessibility) datasets, several genomic annotation contexts were considered, including introns, exons, intergenic regions, CpG islands, promoters (-1500 bp to +500 bp of transcription start sites), H3K4me3 sites (ENCODE (8) accession ID: ENCFF021JBH, experiment: ENCSR000DUO), and H3K27ac sites (ENCODE (8) accession ID: ENCFF763UAG, experiment: ENCSR919WLM). In addition, unbiased 3kb windows of the whole genome were generated with a step size of 1.5kb.

For DNA methylation, the CpG methylation rate was estimated within each annotation window using the Bayes binomial approximation, as in Smallwood *et. al.* (9).

The GpC methylation, which marks accessible DNA in scNOMe-seq libraries, was introduced *in vitro* using a bacterial GpC methyltransferase enzyme. To remove batch effects resulting from differences in enzymatic activity, data normalization and batch correction were performed as follows. GpC data for the whole genome were aggregated in windows of 500kb in length with 250kb overlap separately for methylated and unmethylated GpC counts for each cell. Per-cell pooled size factors were computed from these 500kb windows using the method of Lun *et. al.* (10) scaling by total library size, as implemented in the single-cell R package scuttle (v1.8.4) (11). Batch scaling factors were estimated from corrected methylated and unmethylated window log counts using the rescaleBatches method from the R package batchelor (v1.14.1) (12). Per cell methylated and unmethylated cell-scaling factors were calculated as the ratio of the batch-corrected sum of counts to the mean sum of counts across cells. Finally, unscaled methylated and unmethylated counts in each cell were independently scaled by the product of the cell pooled size factor and methylated/unmethylated count batch correction factor, respectively. The GpC methylation rate for each annotation window was then computed using normalized batch-corrected counts by Bayesian binomial approximation.

From the overall distribution of counts across the annotation layer for CpG and GpC methylation data, minimum total count thresholds per window of 5 counts (CpG) and 20 counts (GpC) were established and applied to discard windows with unreliable methylation rate estimations.

##### scRNAseq normalisation and batch correction

scRNA-seq libraries from HL-60 scNMT-seq data were filtered to remove lowly expressed genes, requiring at least five counts in 10% of the cells. Normalization and variance stabilization were performed by the scTransform method (13) and batch correction by anchor-based integration using the R package Seurat (v4.2.0) (14). First, the batches were independently normalized using scTransform. Then, the top 5,000 most variable features that were in common across batches were identified and used to determine integration anchors. These features and anchors were then used for integration, thereby retaining those 5,000 commonly variable features. Finally, a sparse RNA-seq matrix was utilized for some analyses, whereby gene imputation calculations, performed by Seurat during integration, were ignored and removed by reintroducing ‘NAs’ in place of genes with originally ‘missing data’ (zeros). All downstream analyses only considered autosomal genes (Chr1-22).

##### Colony RNA-seq quality control, normalization and batch correction

Samples were excluded if they had less than 35% genes measured, or less than 70% reads in exons for HL-60 and MOLM-13 samples, or less than 65% reads in exons for MV-4-11 samples. RNA-seq data from single colonies were filtered to remove lowly expressed genes, requiring at least five counts in three samples. For each cell line, normalization was performed by ‘scuttle::logNormCounts()’ (v1.6.2) and batch corrected using mutual nearest neighbors method by ‘batchelor::mnnCorrect()’ (v1.12.3) with default parameters. Downstream analyses only considered autosomal genes (Chr1-22).

##### Correlation analysis in single-colony data

Pearson correlations comparing gene expression to mean global methylation in DAC HL-60 colonies (Fig. S6) were performed using ‘cor.test()’ and underwent Benjamini–Hochberg false discovery rate adjustment using ‘p.adjust(method=”BH”)’. Gene clustering and heatmap visualization were performed for significantly correlated genes (p.adj ≤ 0.05 & cor.value.estimate ≤ -0.4 or cor.value.estimate ≥ 0.4). The average expression of each gene was calculated for each treatment group, with DAC split into high (≥75%) and low (<75%) global methylation groups. The R package “pheatmap” (v1.0.12) was used to plot the mean centered treatment group average expression levels with rows aggregated into 4 ‘kmeans_k’ clusters. The genes from each ‘Kmeans_k’ cluster were extracted and underwent GO ORA for biological process individually using 'enrichGO()’ with fdr adjustment, and the results are displayed as tree plots.

##### Consideration of ARRIVE guidelines relating to animal experiments

###### Animal housing

NSG mice were kept at the Bioresources Facility (Callaghan, NSW, Australia) at 22 ± 2°C, with water and chow *ad libitum*, under a 12:12 h light and dark photoperiod, and housed in ventilated cages in a pathogen-free environment.

###### Group size

G*Power 3.1 (15) was used to perform calculations on sample size, effect size, and statistical power. The minimal significance (α) and statistical power (1-β) were set at 0.05 and 0.80, respectively. Calculations were carried out for two groups by using Student’s t-distribution.

###### Randomisation and blinding

For the cell line-derived xenograft model (MOLM-13-*luc)*, once the luminescence signal was detected (day 6 post-engraftment), mice were randomly allocated to treatment groups. The average BLI of each group was calculated to ensure that tumour burden was equivalent between treatments. Mice with low BLI (negative signal at day 6 post-engraftment) were excluded from the study. For the patient-derived xenograft model (AML-16), once around 1% of leukemia burden was detected in the blood of mice (~3.5 weeks post-engraftment), mice were randomly allocated to treatment groups. The average % of leukemia burden of each group was calculated to ensure that burden was equivalent between treatment groups.

###### Animal monitoring

Following xenotransplantation, animals were monitored daily for clinical and behavioural changes, and body weight was recorded twice a week with a digital balance. Ethical endpoint was defined according to a monitoring checklist (inhibited activity, lethargy, moribund animal, loss of ≥20% of initial body weight, ≥25% leukemia burden – for AML-16 model). All animals were euthanised by carbon dioxide asphyxiation at ethical endpoint.

#### Supplementary Figures

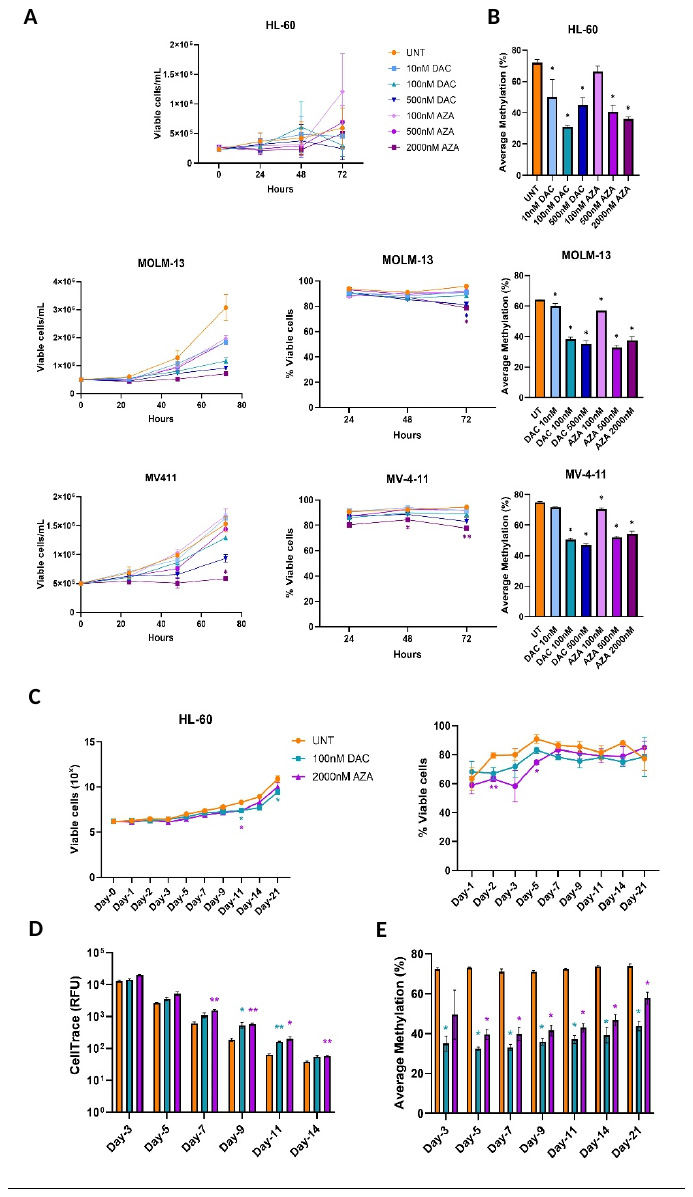

**Fig. S1: Low-dose HMA treatment reduces DNA methylation without acute cytotoxicity in AML cell lines.** AML cell lines (HL-60, MOLM-13, and MV-4-11) were treated every 24 hours for a total of 72 hours, with various doses of decitabine (DAC: 10, 100, 500nM) and azacytidine (AZA: 100, 500, 2000nM), or untreated (UNT) in suspension culture. **A)** Cell growth and viability was measured throughout the dose response experiment using trypan blue exclusion assay (viable cells/mL and % viable cells). **B)** Following 72 hours of HMA treatment, average DNA methylation levels (%) were assessed using Post-bisulphite Adapter Tagging (PBAT). Time-course experiment showing: **C)** cell viability (total viable cells, left; % viable cells, right), **D)** cell division (CellTrace fluorescence), and **E)** average methylation (PBAT) at various time points following treatment with 100nM DAC and 2000nM AZA in HL-60 cells. All data shown as mean +/- SEM, *n=3* experiments. Statistical analysis (A) was performed using two-way repeated measures (RM) ANOVA with Dunnett's multiple comparisons test (p<0.04*, p<0.002**), compared to respective UNT within each time point. Statistical analysis (B) was performed using ordinary one-way ANOVA with Dunnett's multiple comparisons test (*p* < 0.03*), compared to UNT. Statistical analysis (C - E) was performed using two-way repeated measures (RM) ANOVA with Dunnett's multiple comparisons test (*p* < 0.04*, *p* < 0.008**), compared to respective UNT within each time point. Related to Figures 1 and 3.

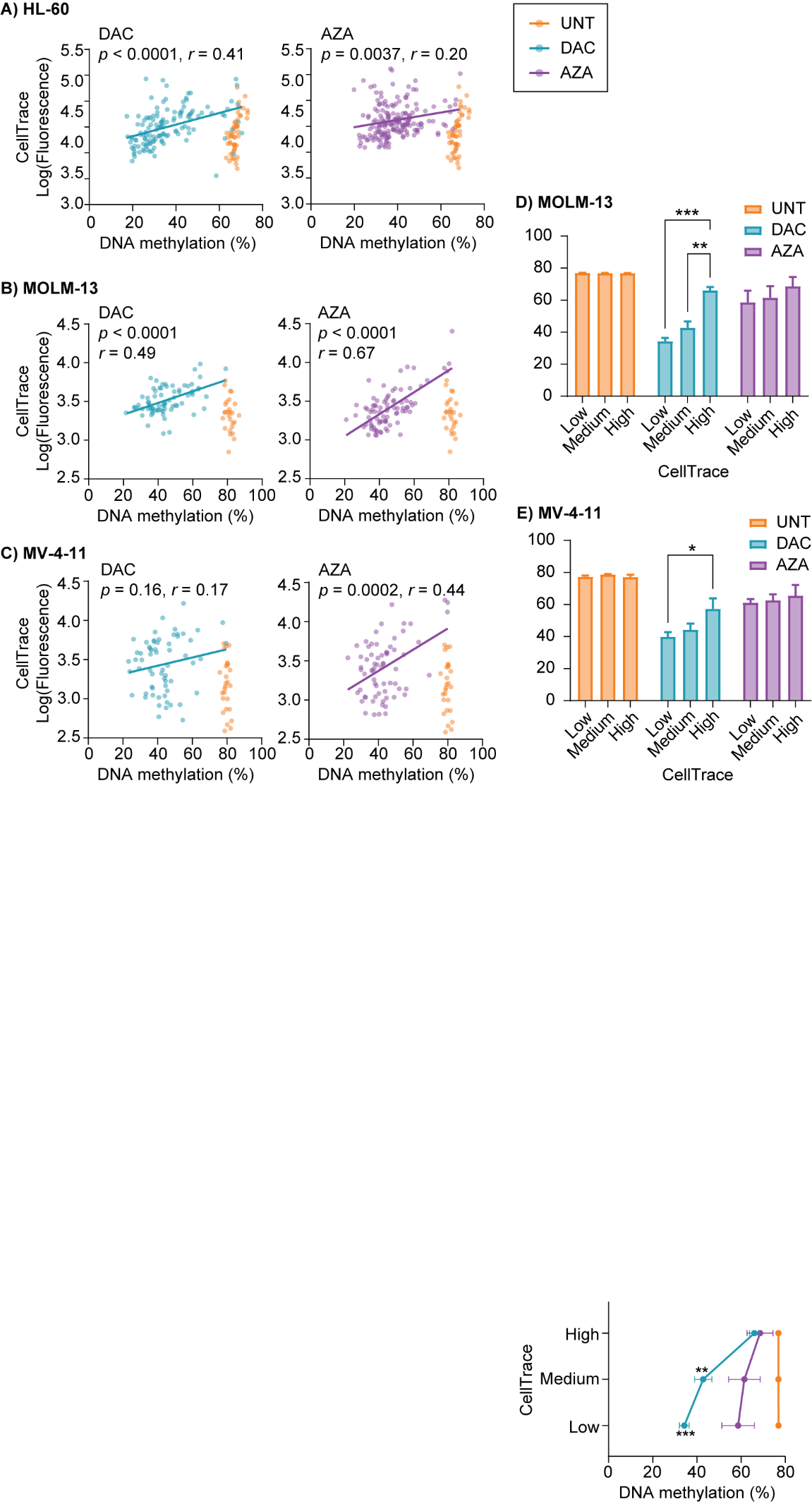

**Fig. S2: Slowly dividing AML cells retain high levels of DNA methylation during HMA treatment.** HL-60, MOLM-13 and MV-4-11 cells were labelled with CellTrace and treated with decitabine (DAC; 100nM) or azacytidine (AZA; HL-60: 2000nM, MOLM-13 and MV-4-11: 500nM) every 24h for 72h. **A - C)** Data from Figure 1B are replotted with the addition of untreated cells from Figure 1A (UNT, orange), and an additional 138 scBS-seq libraries from HL-60 cells (UNT = 16, DAC n=38, AZA n = 84). Linear regressions for DAC (cyan, left) and AZA (purple, right) groups are shown with F-test *p*-values, and Pearson correlation coefficients (*r*). **D - E)** Populations of CellTrace-high, -medium, and -low MOLM-13 and MV-4-11 cells were collected by FACS and TEM-seq analysis of DNA methylation at SINE Alu sites was performed. Data are shown as mean and range for duplicate experiments. Statistical analysis was performed using ordinary two-way ANOVA with Dunnett’s multiple comparisons test: * < *p* < 0.05, ** *p* < 0.01, *** *p* = 0.001 vs. CellTrace-high. Related to Figure 1.

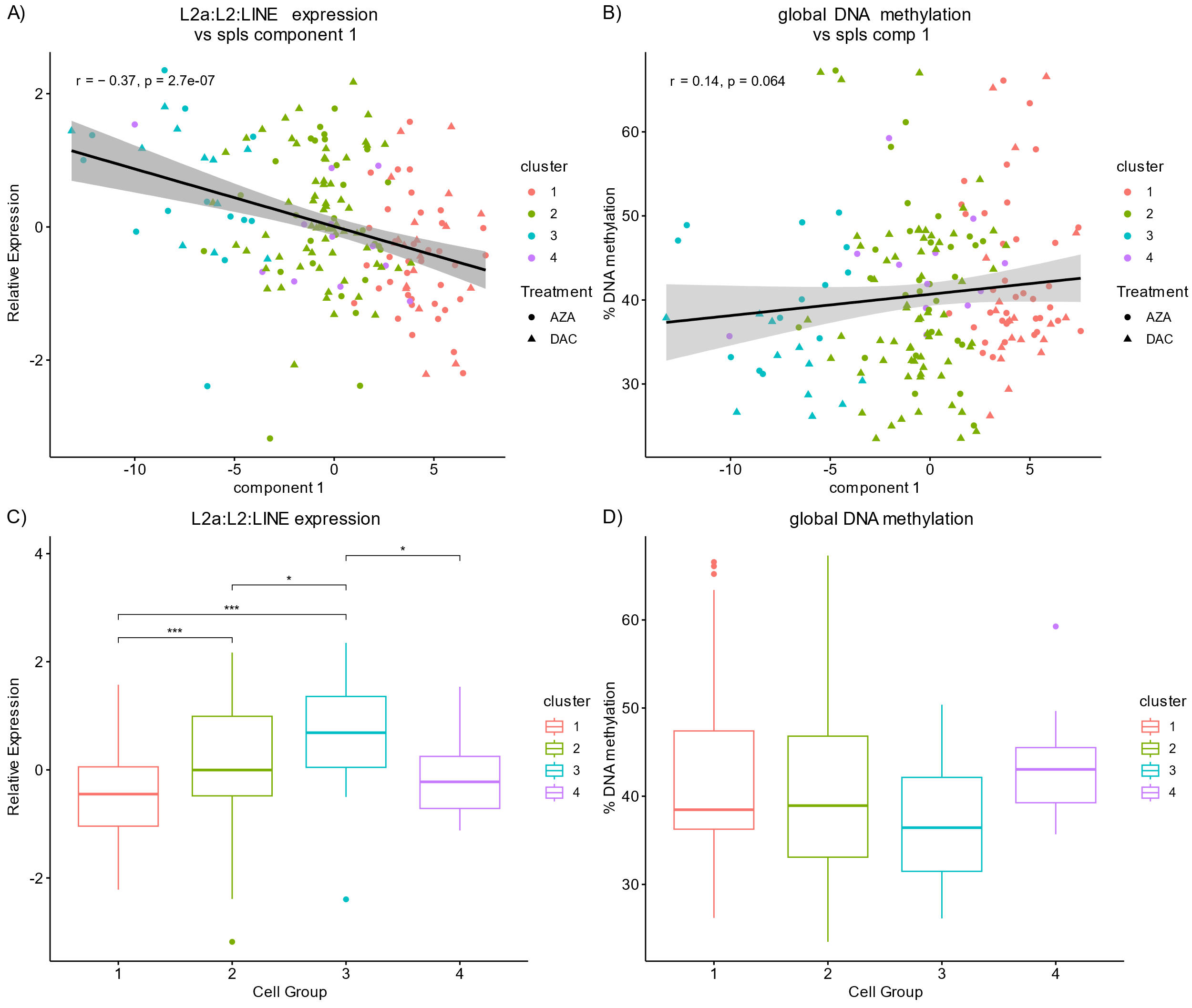

**Fig. S3**: **LINE:L2a expression and DNA methylation correlations with sPLS component 1 and comparisons between cell groups.** **A)** Expression of L2a and **B)** global DNA methylation plotted against sPLS component 1 values for HMA treated cells. Fitted linear models and statistical testing by spearman’s rank correlation are included. **C)** L2a expression and **D)** global DNA methylation are compared between cell groups identified from sPLS analysis (Fig. 2D). Boxes depict the interquartile range (IQR) with median. Whiskers extend to the highest and lowest data points within 1.5 x IQR of the first and third quartile. Significance was determined by Kruskal Wallis testing and post-hoc analysis by pair-wise Wilcoxon rank sum testing with Benjamini-Hochberg correction: * p ≤ 0.05, ** p ≤ 0.01, ***, p ≤ 0.001.

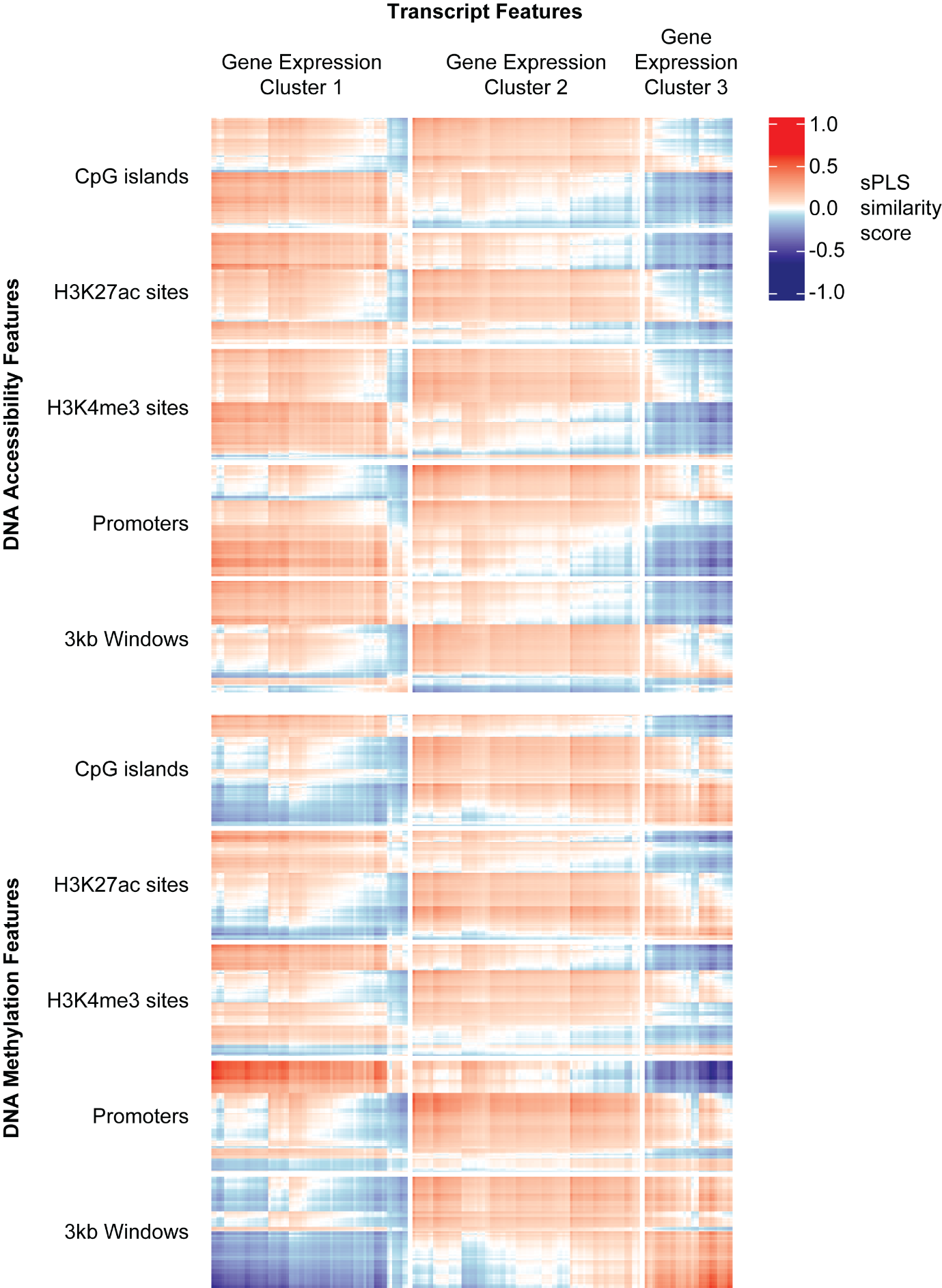

**Fig. S4: Transcript and epigenetic features selected by sPLS (r**elated to Figure 2). Similarity score heatmap computed by mixomics::circosPlot() using sPLS shown in figure 2.

**B)**

**A)**

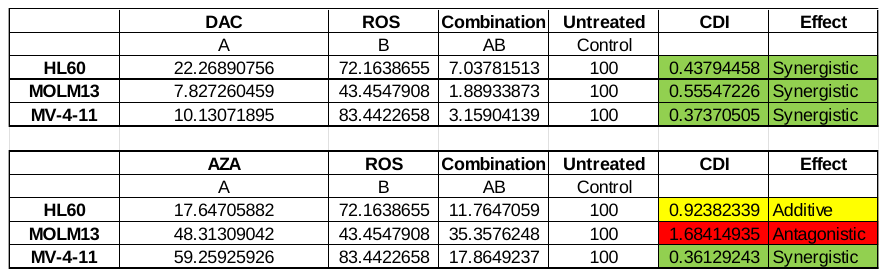

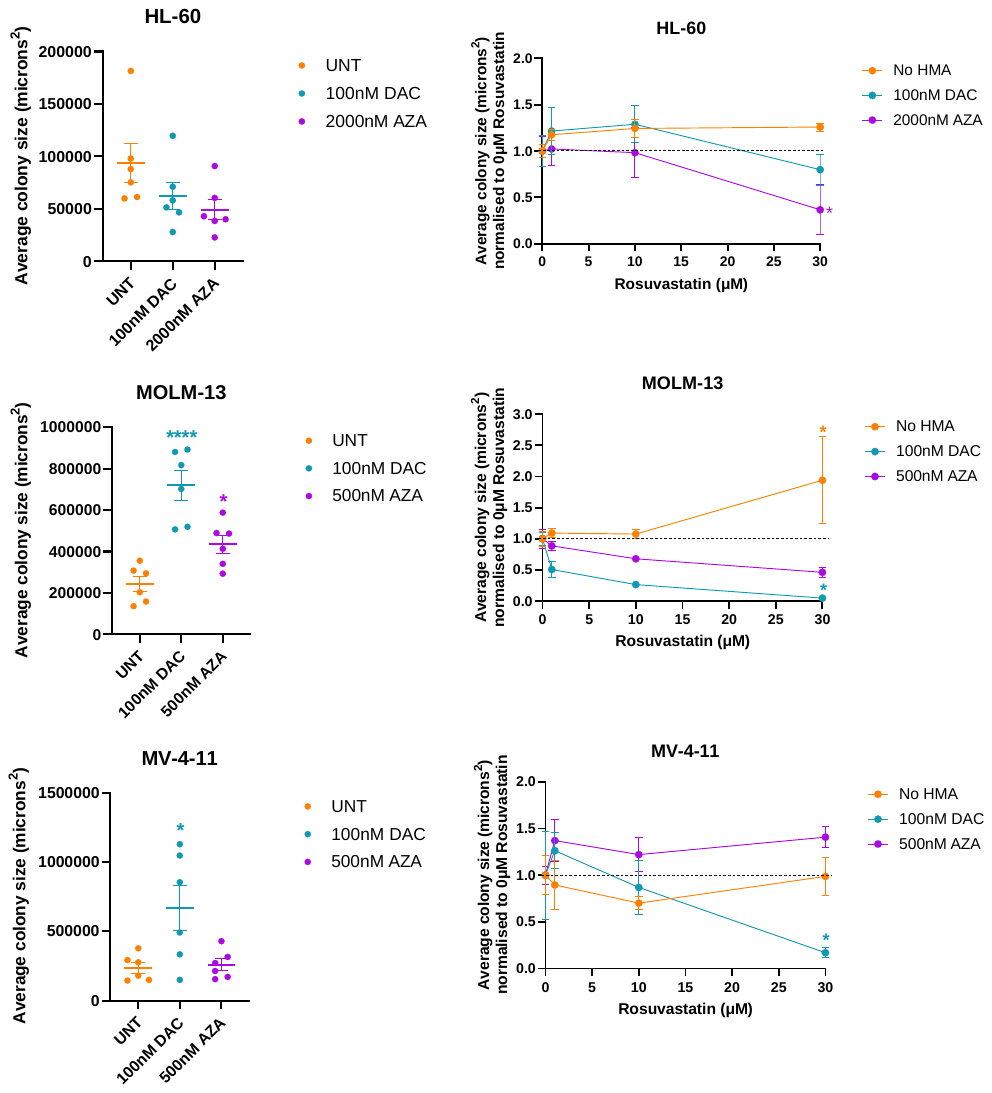

**C)**

**Fig. S5: AML colonies formed after HMA treatment and HMA/statin co-treatment differ in size.** **A)** Average size of AML cell colonies (microns^2^) formed following treatment with DAC (cyan) or AZA (purple) for 72 hours prior to colony-forming assays. UNT = untreated, *n = 6*. **B)** Average size of AML cell colonies following treatment with no HMA, DAC or AZA for 72 hours prior to colony-forming assays with various doses of rosuvastatin (0, 1, 10, 30µM). For each HMA treatment, colony sizes are normalised to the 0µM rosuvastatin control, *n = 3*. Statistical analysis using Ordinary one-way ANOVA with Dunnett's multiple comparisons test (*p* < 0.05) for panel A (*p* < 0.0001****, *p* < 0.03*) compared to UNT, and Ordinary two-way ANOVA with Dunnett's multiple comparisons test (*p* < 0.05) for panel B (*p* < 0.04*) compared to corresponding 0µM rosuvastatin control. All data is shown as mean +/- SEM. Related to Figures 3 and 5. **C)** Colony counts from Figure 5B were normalised to untreated colony counts and the coefficient of drug interaction (CDI) was calculated using the following formula: CDI = AB / (A x B). CDI values of 1, <1 or >1 express additive, synergistic or antagonistic effects, respectively. A CDI value < 0.7 demonstrates a strong synergism of the drug combination (HMA + ROS). DAC (100nM) combined with ROS (30µM) showed synergism in all three AML cell lines with respect to inhibition of self-renewal capacity (colony formation), however, synergism was only seen in MV-4-11 cells when AZA (500nM) was combined with ROS (30µM).

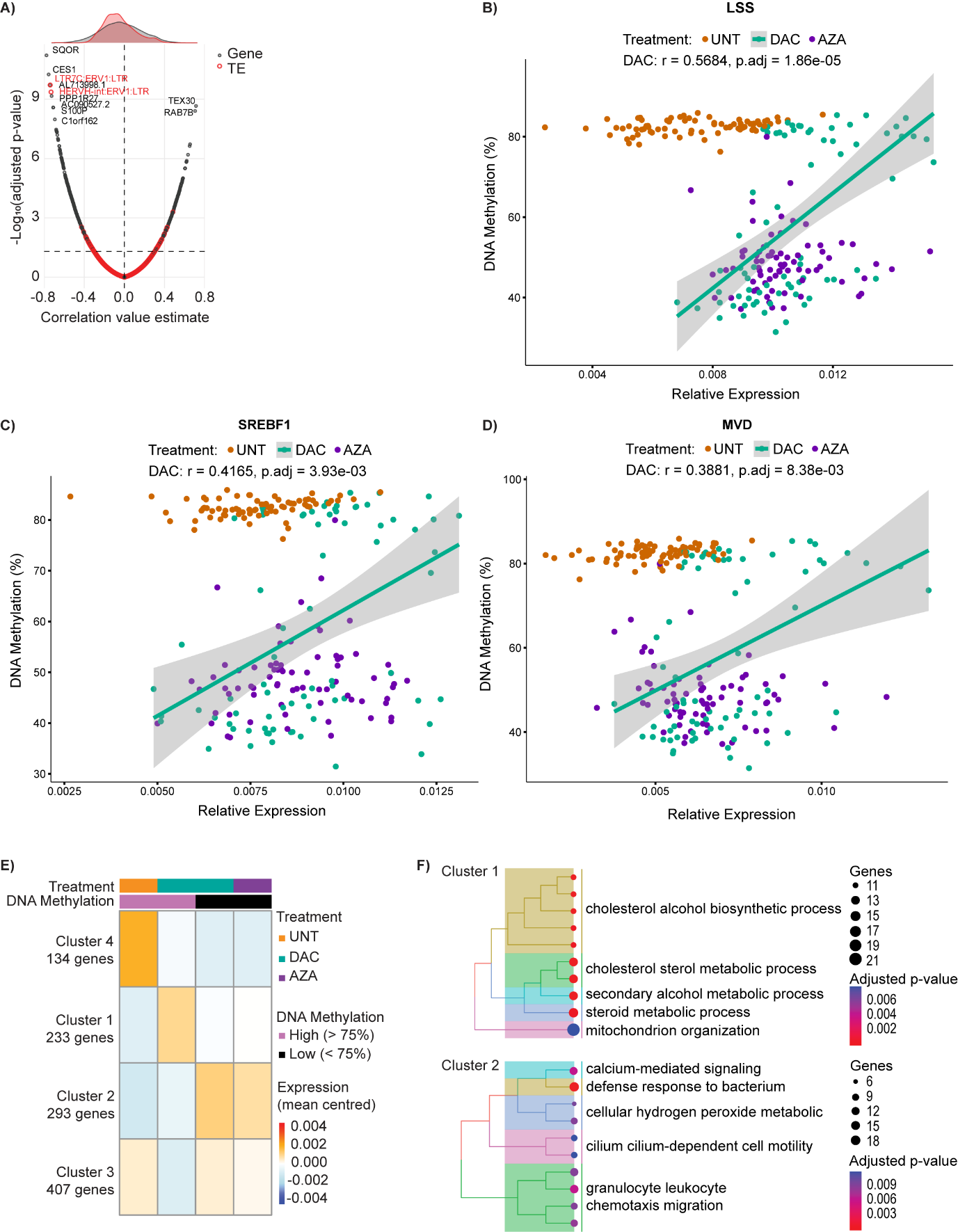

**Fig. S6: Correlations between gene expression and global DNA methylation in HL-60 colonies formed after decitabine (DAC) treatment. A)** Volcano plot showing the Pearson correlation coefficient and adjusted *p*-value for correlations between gene or transposable element (TE) expression and global DNA methylation levels from HL-60 colonies derived following DAC treatment. The upper density plot shows a bias toward negative correlations, especially between global DNA methylation level and TE expression (red). **B) to D)** Scatter plots display representative correlations for three genes from ‘cholesterol biosynthetic process’ (GO:0006695): *LSS* (B), *SREBF1* (C), *MVD* (D). Linear model regression lines for DAC-treated cells were plotted by R displaying standard error intervals. Correlation coefficients (r) and adjusted p-values are from the correlation analysis in Table S10. Data from untreated (UNT) and azacytidine (AZA) samples are also shown. **E)** Simplified heatmap of k-means clustering for the 1,067 genes with significant correlations to global DNA methylation level (adjusted *p*-value ≤ 0.05 and 0.4 ≤ correlation estimate ≤ 0.4, from A). **F)** Summarised tree plots displaying GO terms with significant (adjusted *p*-value < 0.05) over-representation in clusters 1 and 2 (from E).

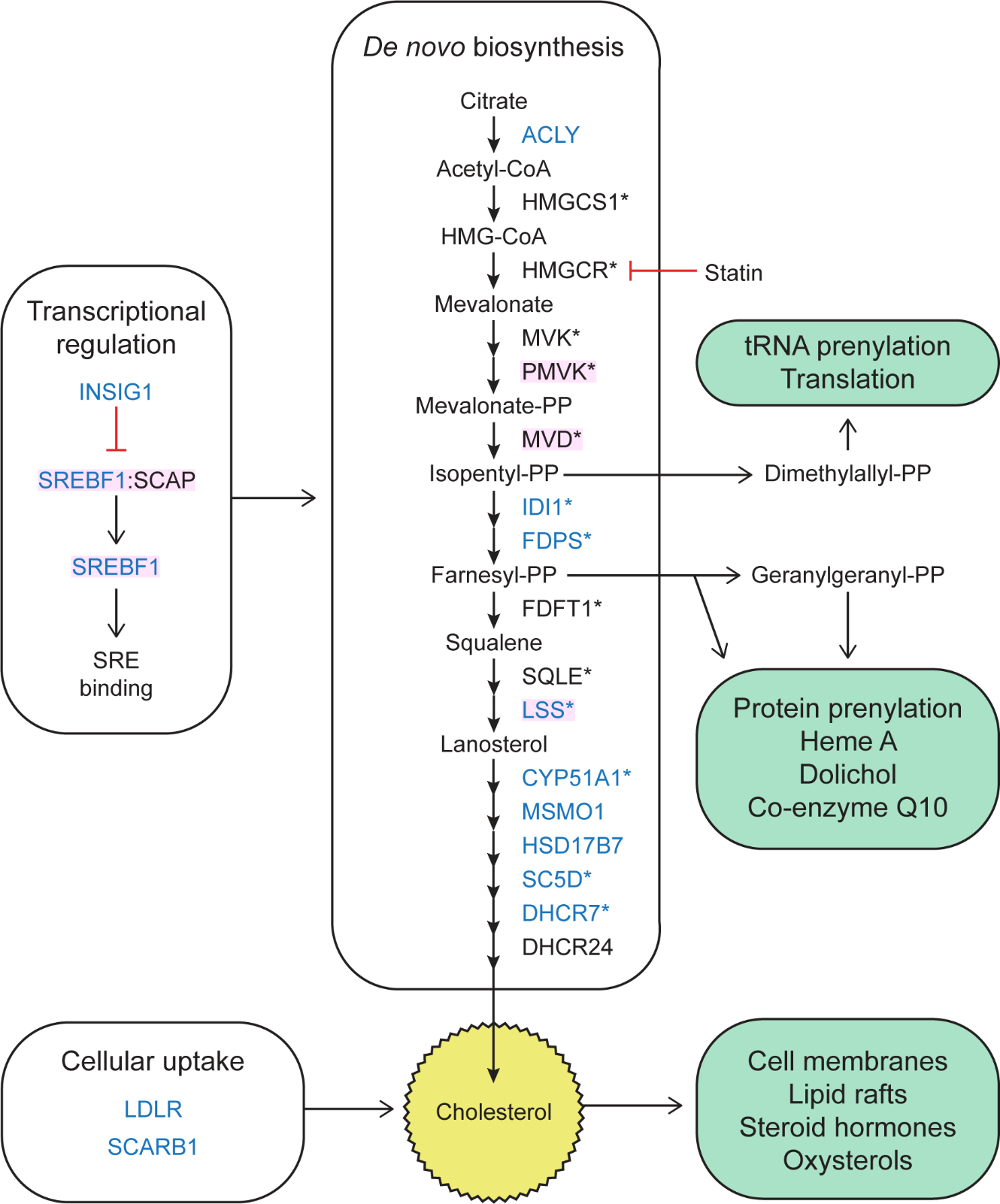
**­­**

**Fig. S7: Genes involved in cholesterol biosynthesis.** Schematic illustration showing the contribution of selected genes to cholesterol regulation. Pink shading indicates genes that were significantly increased (Pair-wise Wilcoxon test with Benjamini-Hochberg correction) in all cell lines by both DAC and AZA (Fig. 5A). Blue symbols indicate genes with positive correlations between expression and global DNA methylation among DAC-treated HL-60 colonies (cluster 1, Supplementary Fig. 6E). *SREBF1* encodes the SREBP1 transcription factor, which binds to sterol regulatory elements (SRE) to control the expression of target genes. Genes with promoter SREBP1 binding (according to Reactome pathway #R-HSA-2426168) are marked with an asterisk. Statins inhibit the HMGCR enzyme. Green boxes list downstream functions of cholesterol and other metabolites produced during cholesterol biosynthesis. Related to Figure 5.

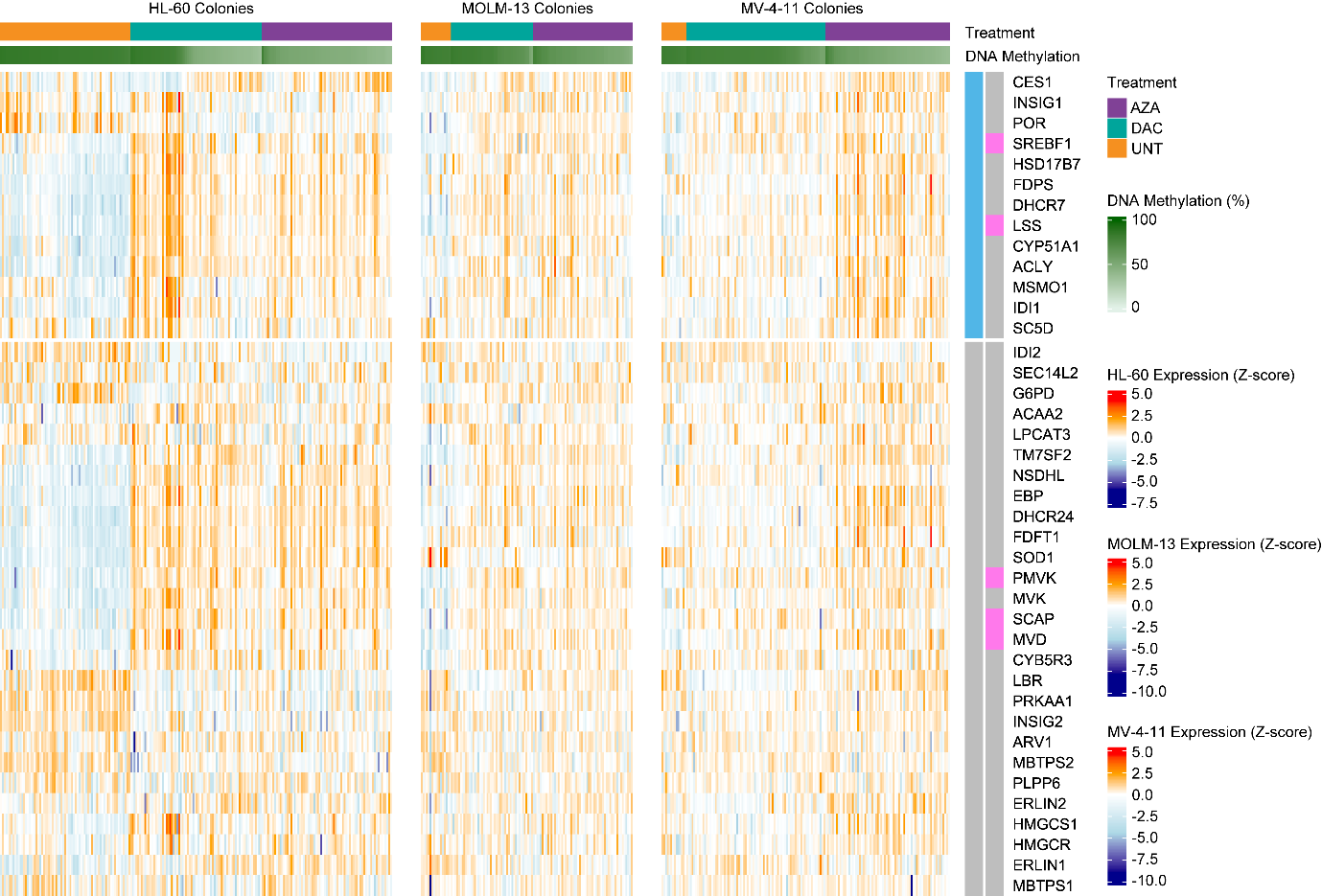

**Fig. S8: Expression of genes from ‘cholesterol biosynthesis process’ (GO:0006695) in colonies from all 3 cell lines.** Genes with positive correlations between expression and global DNA methylation among DAC-treated HL-60 colonies (cluster 1, Fig. S6E) are shown at the top (blue side bar). Genes that were significantly increased in all cell lines by both DAC and AZA (Pair-wise Wilcoxon test with Benjamini-Hochberg correction, Figure 5A) are also indicated (pink side bar). Related to Figure 5.

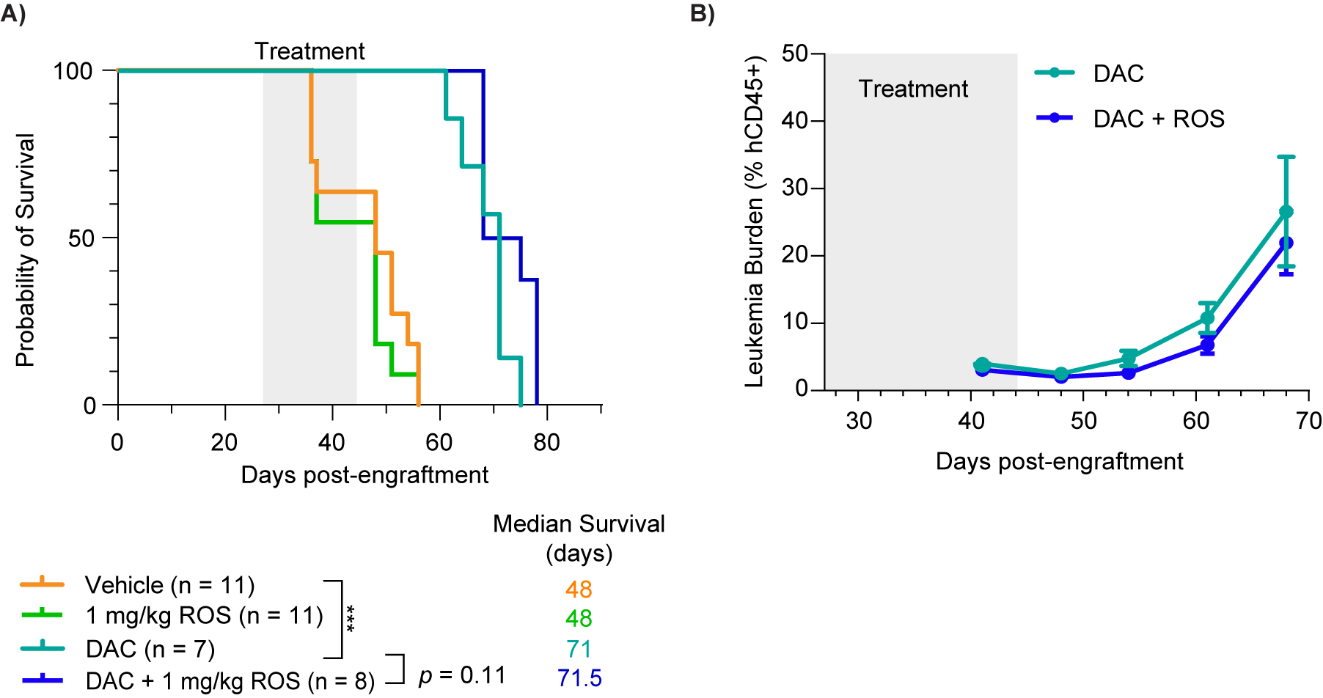

**Fig. S9: Leukemia burden and survival in AML xenografts.** AML-16 cells were engrafted into NSG mice and mice were treated with vehicle control, rosuvastatin (1 mg/kg/day), DAC (0.2mg/kg/day), or DAC (0.2mg/kg/day) + rosuvastatin (1 mg/kg/day) on a treatment schedule of ‘5 days on, 2 days off’ for the first cycle, followed by two times per week for the remaining two cycles. **A)** Survival analysis was performed using Kaplan-Meier analysis followed by the Log-rank (Mantel-Cox) test in Graphpad, and an adjusted *p*-value of < 0.05 was considered statistically significant. **B)** Leukemia burden (% human CD45^+^ cells) post-engraftment and following the cessation of treatment at various timepoints. Mann-Whitney test (unpaired, non-parametric, two-tailed t-test) was used to test for statistical significance, with a p-value cut-off of 0.05 for significance.

#### Supplementary Tables

**Table S1: Characteristics of AML cells used in this study.** Information on HL-60, MOLM-13 and MV-4-11 cell lines were gathered from DSMZ (Sex, Age, FAB Subtype) and Cellosaurus (Mutated Genes and Gene Fusions). Data for AML-16 are taken from Lee *et al* (16).

| **Cells** | **Sex** | **Age** | **FAB Subtype** | **Mutated Genes** | **Gene Fusions** |
| --- | --- | --- | --- | --- | --- |
| HL-60 | F | 35 | M2 | TP53; CDKN2a; NRAS | None |
| MV-4-11 | M | 10 | M5 | FLT3 | MLL-rearranged (KMT2A-AFF1) |
| MOLM-13 | M | 20 | M5a | FLT3 | MLL-rearranged (KMT2A-MLLT3) |
| AML-16 | F | 61 | M4 | FLT3; IDH2; NPM1; WT1 | None |

**Table S7: Summary of region-wise correlations.** Pearson correlations were computed between gene expression and DNA methylation (top) or accessibility (bottom) of nearby loci. The total number of correlations computed as well as the % negative (*p* < 0.05, cor < 0, red) and % positive (*p* < 0.05, cor > 0, green) are shown for all genes, and for each gene expression cluster from the sPLS model. Related to Figure 2.

|  |  | **All Genes** | | | **Cluster 1 Genes** | | | **Cluster 2 Genes** | | | **Cluster 3 Genes** | | |
| --- | --- | --- | --- | --- | --- | --- | --- | --- | --- | --- | --- | --- | --- |
|  |  | Total | Negative | Positive | Total | Negative | Positive | Total | Negative | Positive | Total | Negative | Positive |
| **DNA Methylation** | H3K4me3 sites | 6177 | 1.7% | 3.2% | 87 | 1.1% | 2.3% | 112 | 1.8% | 3.6% | 66 | 1.5% | 4.5% |
|  | H3K27ac sites | 6794 | 1.7% | 3.0% | 130 | 0.8% | 3.1% | 134 | 2.2% | 2.2% | 55 | 0.0% | 3.6% |
|  | CpG islands | 4715 | 1.9% | 3.6% | 47 | 2.1% | 4.3% | 82 | 1.2% | 2.4% | 50 | 2.0% | 6.0% |
|  | Promoters | 31645 | 1.7% | 3.3% | 412 | 6.1% | 1.5% | 607 | 0.8% | 3.3% | 271 | 1.8% | 3.7% |
|  | Promoter-proximal 3kb windows | 99926 | 2.5% | 2.9% | 1539 | 4.6% | 2.3% | 1658 | 1.2% | 5.5% | 779 | 1.6% | 4.8% |
| **DNA Accessibility** | H3K4me3 sites | 6177 | 2.5% | 2.6% | 87 | 1.0% | 8.2% | 112 | 1.7% | 3.4% | 66 | 2.9% | 0.0% |
|  | H3K27ac sites | 6794 | 2.6% | 2.5% | 130 | 2.5% | 6.7% | 134 | 1.4% | 2.7% | 55 | 6.7% | 0.0% |
|  | CpG islands | 4715 | 2.4% | 2.9% | 47 | 4.3% | 6.4% | 82 | 0.0% | 4.9% | 50 | 2.0% | 2.0% |
|  | Promoters | 31645 | 2.5% | 2.8% | 412 | 1.9% | 7.3% | 607 | 0.4% | 4.6% | 271 | 2.1% | 1.4% |
|  | Promoter-proximal 3kb windows | 99926 | 2.3% | 3.0% | 1539 | 1.7% | 4.4% | 1658 | 2.6% | 4.2% | 779 | 3.5% | 2.3% |

**Table S13: Statistical analysis related to Figure 6A.** The exact p-values are shown for comparisons between treatment groups with significant differences highlighted by green shading.

|  | **DAC (0.2 mg/kg)** | **DAC (0.2 mg/kg) + ROS (1 mg/kg)** | **DAC (0.2 mg/kg) + ROS (10 mg/kg)** | **DAC (0.2 mg/kg) + ROS (40 mg/kg)** |
| --- | --- | --- | --- | --- |
| **VEH** | 0.0011 | <0.0001 | <0.0001 | <0.0001 |
| **DAC (0.2 mg/kg)** |  | 0.0012 | <0.0001 | 0.0026 |

**Table S14: Statistical analysis related to Figure 6B.** The exact p-values are shown for comparisons between treatment groups with significant differences highlighted by green shading.

|  | **DAC (0.2 mg/kg)** | **ROS (1 mg/kg)** | **DAC (0.2 mg/kg) + ROS (1 mg/kg)** |
| --- | --- | --- | --- |
| **VEH** | 0.0457 | 0.0064 | 0.0002 |
| **DAC (0.2 mg/kg)** |  | <0.0001 | 0.0053 |
| **ROS (1 mg/kg)** |  |  | <0.0001 |

**Table S15: Statistical analysis related to Figure 6C and Fig. S9C.** The exact p-values are shown for comparisons between treatment groups with significant differences highlighted by green shading.

|  | **DAC (0.2 mg/kg)** | **AZA (1 mg/kg)** | **ROS (1 mg/kg)** | **DAC (0.2 mg/kg) + ROS (1 mg/kg)** | **AZA (1 mg/kg) + ROS (1 mg/kg)** |
| --- | --- | --- | --- | --- | --- |
| **VEH** | <0.0001 | 0.0002 | 0.3684 | <0.0001 | <0.0001 |
| **DAC (0.2 mg/kg)** |  | 0.9361 | <0.0001 | 0.1118 | 0.0068 |
| **AZA (1 mg/kg)** |  |  | 0.0001 | 0.1262 | 0.0086 |
| **ROS (1 mg/kg)** |  |  |  | <0.0001 | <0.0001 |
